# Recurrent issues with deep neural network models of visual recognition

**DOI:** 10.1101/2024.04.02.587669

**Authors:** Tim Maniquet, Hans Op de Beeck, Andrea Ivan Costantino

**Affiliations:** KU Leuven, Leuven, Belgium

## Abstract

Object recognition requires flexible and robust information processing, especially in view of the challenges posed by naturalistic visual settings. The ventral stream in visual cortex is provided with this robustness by its recurrent connectivity. Recurrent deep neural networks (DNNs) have recently emerged as promising models of the ventral stream, surpassing feedforward DNNs in the ability to account for brain representations. In this study, we asked whether recurrent DNNs could also better account for human behaviour during visual recognition. We assembled a stimulus set that included manipulations that are often associated with recurrent processing in the literature, like occlusion, partial viewing, clutter, and spatial phase scrambling. We obtained a benchmark dataset from human participants performing a categorisation task on this stimulus set. By applying a wide range of model architectures to the same task, we uncovered a nuanced relationship between recurrence, model size, and performance. While recurrent models reach higher performance than their feedforward counterpart, we could not dissociate this improvement from that obtained by increasing model size. We found consistency between humans and models patterns of difficulty across the visual manipulations, but this was not modulated in an obvious way by the specific type of recurrence or size added to the model. Finally, depth/size rather than recurrence makes model confusion patterns more human-like. Contrary to previous assumptions, our findings challenge the notion that recurrent models are better models of human recognition behaviour than feedforward models, and emphasise the complexity of incorporating recurrence into computational models.

## Introduction

Visual recognition is at the core of human cognition, and an impressive feature of our brain. Understanding the visual world seems effortless, which hides the computational difficulty of real-world object recognition. The visual complexity of scenes from everyday life calls for flexible and robust processing in order to decode the identity of objects in the environment [Kravitz et al., 2013, Grill-Spector and Weiner, 2014, Bracci and Op de Beeck, 2023]. This feat is accomplished seamlessly by the ventral stream in visual cortex, whose connectivity allows for such extraordinary abilities. While it is often modelled as a bottom-up network, every feedforward connection it contains is paralleled by one or several non-feedforward connections ([Van Essen et al., 1986, Felleman and Van Essen, 1991, Lamme et al., 1998, Ungerleider et al., 2008, Baizer et al., 1991]). It is hence better understood as a highly recurrent network.

Recurrence in a system allows for the dynamic processing of information. Visual recognition, although fast and largely bottom-up ([DiCarlo et al., 2012, Riesenhuber and Poggio, 2000, Riesenhuber, 2005]), is therefore a dynamic process. Across cycles of extra-feedforward processing, recurrent activity modulates and integrates visual information. As a result, it is responsible for many perceptual phenomena known to occur particularly in ambiguous, complex visual situations, e.g. stimulus-context modulation, predictive processing and figure-ground segmentation [Pennartz et al., 2019, Wyatte et al., 2014, Lamme and Roelfsema, 2000, Kreiman and Serre, 2020]. Through the processing flexibility that it confers, recurrence is hence largely implicated in the presence of non-canonical, non-trivial contexts. Evidence for this role comes from studies using backward masking. Given the temporal unfolding that is inherently linked with recurrent processing, it is possible to selectively impact it, while leaving initial feedforward processing intact through the use of masking. Several studies have made use of backward masking to manipulate recurrent processing, and found that it selectively impaired visual recognition under challenging conditions, but not under non-challenging ones ([Seijdel et al., 2021, Rajaei et al., 2019, Tang et al., 2018]).

While feedforward processing is well understood, there is no universal account of the exact mechanisms served by recurrence in visual recognition. Across the visual cortex, various specific forms of recurrence have been studied and linked to specific perceptual phenomena. For instance: uncertainty computation ([Ladret et al., 2023]) and pattern completion ([Shin et al., 2023]) by lateral connections within V1, edge detection in cluttered images through lateral processing in V1 ([Self et al., 2013]), prediction about occluded shapes by feedback from the prefrontal cortex (PFC) to V4 ([Choi et al., 2018]), figure-ground modulation by feedback from V4 to V1 ([Klink et al., 2017, Angelucci et al., 2017]), and interactions with spatial frequency transformations by feedback from frontal areas ([Bar et al., 2006, Goddard et al., 2016]). The literature points to a wide range of roles played by a wide range of recurrent connections, operating various computations on incoming sensory information. Overall, recurrent processing seems to determine an important part of visual perception, and of the brain dynamics underlying it.

The importance of recurrence in brain dynamics is also supported by studies using *deep neural networks* (DNNs). Optimised on recognition tasks, DNNs are considered good models of the ventral stream, mimicking its hierarchical structure and operating classification with human-like accuracy ([Cadieu et al., 2014]). Moreover, DNNs have been found to display representational similarity with the ventral stream ([Yamins et al., 2014, Khaligh-Razavi and Kriegeskorte, 2014]). Strikingly, this similarity increases with recurrent DNNs ([Kar et al., 2019, Kietzmann et al., 2019]), indicating that the representational dynamics of the visual cortex fit better with these of a recurrent network, compared to feedforward-only. The aforementioned studies of backward masking also included DNNs to show that models equipped with recurrent-like features show more resilience to challenging conditions than feedforward-only models.

With a better fit to the brain and improved resilience to image challenges, recurrent DNNs are a promising tool to identify the various roles of recurrence and understand their function in the ventral stream. This is especially important given the observation that recurrent models can be considered time-wrapped equivalents of size-matched feedforward models ([van Bergen and Kriegeskorte, 2020]). A consequence of this observation is that the more biologically plausible, recurrent DNNs fare better on predicting brain activity than size-matched non-recurrent DNNs. However, while recurrent models tend to be better models of the brain than their feedforward-only counterparts, it remains unclear whether they are also better models of human behaviour. This is in part due to a lack of investigations into the pattern of errors and confusions of DNNs. In order to test whether recurrence also improves model fit to human behaviour, such investigations should include a wide range of architectures, and a wide range of visual conditions, to emulate the variety of recurrent connections and of phenomena they are associated with.

The present study aimed at taking a step in this direction by running an extensive test of model performance including different types of DNN recurrence and several image conditions. We combined a variety of image manipulations known to involve recurrence in humans in one large experiment ([Seijdel et al., 2021, O’Reilly et al., 2013, Tang et al., 2018, Rajaei et al., 2019, Goddard et al., 2016]), in which we tested and compared human participants and DNN models. We implemented different types of architecture, including different flavours of recurrence, in a number of DNNs, and compared their performance and error patterns on a classification task with that of humans. Taking advantage of the architectural diversity of our models, we asked whether recurrent models would perform better in the resolution of different kinds of visual complexity, and whether they would better explain the patterns human behaviour, as compared to non-recurrent models.

We found that recurrence in DNNs helped them match with human performance, but not more than added depth. This feeds into the observation that recurrent neural networks can be considered time-wrapped equivalents of size-matched feedforward neural networks ([van Bergen and Kriegeskorte, 2020]). However, our results go a step further, as we also found a decrease in models match with human confusion matrices when equipped with recurrence. Additionally, we were not able to find dissociable patterns of performance across our recurrent models, which indicates that DNNs, as tools to model the visual cortex, are unable to reproduce the variability of phenomena that occur in brain recurrence. Overall, we interpret our results, in light of current developments in the field, as an argument for the development of more elaborate, biologically plausible ways of implementing recurrence in models of visual recognition.

## Materials and methods

### Participants

A total of 231 subjects participated in the study (195 females, age (mean±SEM): 18.7 ± 1.7), of which 13 were excluded due to low performance (average accuracy below 0.7). All participants gave their informed consent before taking part. The protocol was approved by the ethics committee at KU Leuven. The task took place online and lasted approximately 30 minutes.

### Stimuli

Images of objects were selected from real-world scenes of the publicly available online databases MS COCO ([Lin et al., 2014]) and ADE20K ([Zhou et al., 2018]). Eight object categories were included: *person*, *cat*, *bird*, *tree*, *fire hydrant*, *building*, *bus*, and *banana* (see Fig 1). The categories were chosen to cover a diverse range of levels of animacy, real world size and aspect ratio. Stimuli were segmented out from their backgrounds, transformed to a 700*×*700 pixel size, and equalised on their contrast and average luminance levels. 10 exemplar objects were picked for each category, resulting in a stimulus set of 80 images.

**Figure 1:**
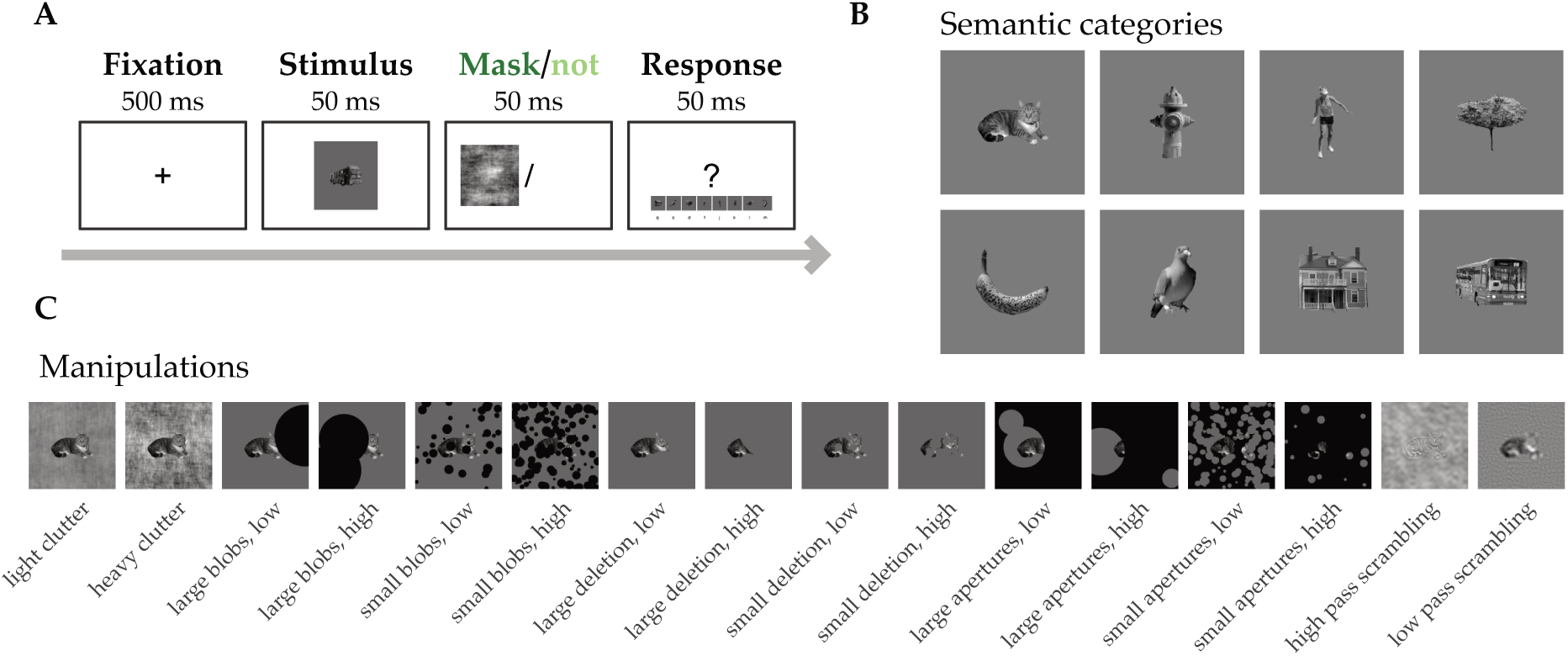
Experimental design & stimuli. (A) Design of the classification task performed by human participants. A 500ms fixation cross (*±* 0-300ms of a randomly long blank interval) was followed by the short presentation of a target stimulus (50ms), which was then either masked (presentation of a phase scrambled pattern for 300ms) or not (blank screen). Finally, a reminder appeared on screen after the mask/blank screen for participants to recall the category-response key mapping. Trials ended upon response or after a maximum of 10 seconds post stimulus. (B) The eight semantic categories included in the stimuli. Top row, from left to right: cat, fire hydrant, person, tree; bottom row, from left to right: banana, bird, building, bus. (C) Challenging manipulations implemented in the stimulus set.

### Challenging manipulations

Object recognition was rendered challenging by applying several types of visual manipulations on the stimulus set (see Fig S1). Each one of the manipulation was selected based on literature demonstrating that it is linked to recurrent processing, as evidenced by psychophysical, neuroimaging, or DNN data. We implemented a total of 16 visual manipulations, each a variation of one of the following image transformations: adding clutter in the background of the object, adding occluding blobs on top of the object, removing parts of the object, adding a full occluder with apertures on top of the object, and phase scrambling the object image. The control condition showed segmented object on a grey background.

We *cluttered* objects by placing cluttered images in their backgrounds. These were obtained by phase scrambling randomly selected natural scenes from the MS COCO dataset, and measuring an index of spatial coherence (SC) and contrast energy (CE) for each image, the combination of which gives a proxy of how cluttered an image is ([Groen et al., 2013]). We defined two levels of clutter by selecting lightly cluttered backgrounds and heavily cluttered backgrounds at each end of the spectrum, with 2 clutter conditions as a result.

We added black, circular *blobs* to occlude 40% (low) or 80% (high) of objects. These blobs were either many small or a few large ones, resulting in 4 blobs conditions.

Parts of objects were *deleted*, with 40% (low) or 80% (high) of objects deleted. The deletion was operated through circular disks, which could also be many small or a few large ones. This resulted in 4 deletion conditions.

A full *occluder* was placed on top of objects, on which apertures were made so that either 40% or 80% of the objects would be hidden. A similar approach with many small or a few large apertures was adopted, with 4 occluder conditions as a result.

*Phase scrambled* images were made by randomising their phase spectrum, adopting an approach similar to that of [Goddard et al., 2016]. Images were Fourier transformed, and their phase spectrum was replaced by random noise, on either side of a spatial frequency threshold of 1.5 cycles per degree (with images assumed to subtend 10 degrees of visual angle on screen). The origin of the phase spectrum was left intact. Images were then transformed back into image space. This resulted in 2 phase scrambling conditions: a low-pass and high-pass phase scrambled version of each image.

### Experimental task

Participants were given a categorisation task, where each image was to be classified in one of the eight possible categories of the stimulus set. All trials had the same timeline (see Fig 1): after a fixation cross, a target image appeared for 50ms. Targets were either followed by an empty screen or by a mask. The instructions asked for an accurate answer that could be registered as soon as the target was presented. All trials ended with the presentation of a reminder of the eight categories and their associated response keys. The latter appeared at the bottom of the screen 200ms after the mask (or empty screen) disappeared.

Masks were phase-scrambled versions of natural scenes taken from MS COCO. Images were randomly selected from the dataset, resized to match target images (700*×*700), and taken to the Fourier space where their phase spectrum was fully replaced by random noise (except for the origin, left intact) before reconstruction into the image space.

In total, 16 experimental conditions and one control made 17 conditions, all of which to be presented with and without masking. The resulting 34 conditions were divided into sub-experiments: occlusion made up 2 sub-experiments (few large occluder disks and many small occluder disks, 6 conditions each). Phase scrambling and clutter were joined together into one other sub-experiment (4 conditions). These three sub-experiments were ran with and without masking, resulting in 6 experiments in total. An extra unmasked control condition was added to all three of the masked experiments to serve as a comparable baseline. All of them were conducted online over the same period, and no participant took part more than once. Data from all six experiments were pooled together, after confirmation that results on the unmasked control, which was similar for all participants, did not differ significantly across experiments (one-way ANOVA on accuracy with experiment as factor, *F* = 0.68*, p* = 0.64).

### Deep neural network architectures

We investigated thirteen distinct DNN models, each varying in architecture and recurrent connections (see Fig 2). These models were specifically chosen to represent a range of complexities, from basic feedforward structures to more sophisticated recurrent networks. We categorised these models into three groups: CORnet models, B models, and VGG models (see Table 1 for more details).

**Figure 2:**
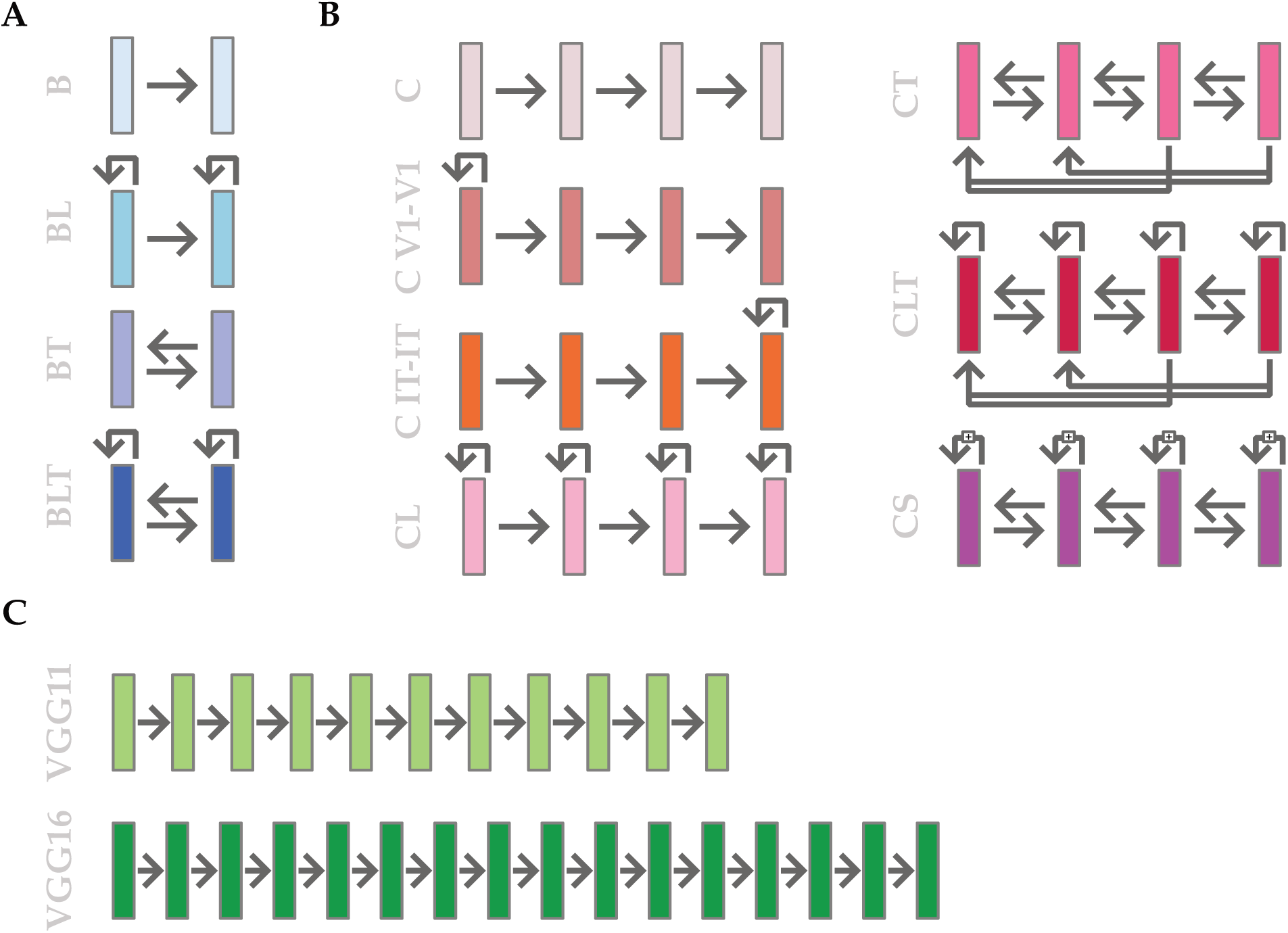
Model architectures. All recurrent and non-recurrent models included in the study. (A) The four available models of the *B* family were used. These models share a baseline 2-layer architecture (B), with the addition of either lateral connections (BL), top-down connections (BT), or both (BLT). (B) Models from the *CORnet* family (here called *C* for facility) including three readily available and four custom-made models. From [Kubilius et al., 2018], the base FF model *CORnet Z* (C), the lateral connections-equipped *CORnet RT* (CL) and the highest-performing *CORnet S* (CS). Custom-made versions of these architectures include full top-down connections (CT), full top-down and lateral connections (CLT), specific lateral connections in layers V1 (C V1-V1) and IT (C IT-IT). (C) *VGG* models were used as feedforward controls to the recurrent models. We used the state-of-the-art VGG16 model, as well as a custom-made, smaller VGG11 to better match the size of our recurrent models. Rightward arrows represent feedforward connections. Backward arrows above layers represent lateral connections. Leftward arrows and arrows below layers represent feedback connections. The arrows of CS represent the ResNet-inspired residual block structures.

#### CORnet Models

From the CORnet family, we included the foundational models CORnet Z, CORnet RT, and CORnet S ([Kubilius et al., 2018, Kubilius et al., 2019]). While CORnet Z uses a simple feed-forward architecture, CORnet RT enhances this design by integrating biologically-inspired additive recurrent mechanisms within each block. CORnet S further advances this architecture by adding additional convolutional layers in a bottleneck ResNet-like block structure, thereby enriching its internal dynamics and significantly increasing its parameter count relative to its simpler counterparts. For better readability, CORnet is hereafter abbreviated *C*, and models are branded after the recurrent element of connectivity they contain (*L* for *lateral* connections, *T* for *top-down* connections). We therefore selected *C* (CORnet Z), *CL* (CORnet RT) and *CS* (CORnet S). We custom-extended the C series for this research, by adding the following: C V1-V1, featuring within-layer connections in its initial convolutional layer; C IT-IT, implementing similar within-layer connections but in its terminal convolutional layer; CT, designed with top-down connections extending from higher to lower layers; and CLT, a specialized version of CL augmented with top-down connections analogous to those in CT.

#### B Models

B models ([Spoerer et al., 2017]) include B, BL, BT, and BLT models. BL is characterized by lateral connections, BT by top-down connections, while the BLT model combines both lateral and top-down mechanisms. These models were used in their original implementations, see ([Spoerer et al., 2017]) for a more detailed description.

#### VGG Models

The VGG models, specifically VGG11 and VGG16 [Simonyan and Zisserman, 2015], served as the feed-forward control group. Their architecture, lacking feedback mechanisms, is deeper, providing a baseline comparison against recurrent models. VGG11 is a custom implementation of VGG16 featuring only 11 trainable layers and a smaller decoding head with a single linear layer. This modification was made to create a VGG16-like very deep feedforward network, but with a number of learnable parameters more comparable to the recurrent models.

### Training Regimen

Our model training process was divided into two distinct phases: a training phase on the ImageNet dataset, followed by a phase of fine-tuning on the specific 8 categories contained in our dataset.

#### ImageNet Training

Models lacking pre-existing ImageNet-trained weights or featuring custom recurrent mechanisms (i.e. C V1-V1, C IT-IT, CT, CLT, VGG11) were trained in-house. This phase involved a single cycle of 25 epochs, using a fixed seed of 42 and a batch size of 256. A one-cycle learning rate policy [Smith and Topin, 2017], as implemented in the *fastai* [Howard et al., 2018] library, was employed. The maximum learning rate for the `fit_one_cycle` algorithm was set by multiplying the optimal rate, determined using *fastai*’s implementation of the learning rate finder [Smith, 2015], by a factor of 10. The training was performed on the ILSVRC2012 challenge subset of the ImageNet dataset [Russakovsky et al., 2015], which includes 1,000 classes and approximately 1.2 million images for training and an additional 50,000 for validation purposes. During the training phase, each image was normalized using the mean and standard deviation statistics from ImageNet, followed by a random crop with a minimum scale of 0.35 which was then resized to 224 x 224 pixels. In the validation phase, images were directly normalized and resized to 224 x 224 pixels, omitting the cropping process.

#### Fine-Tuning

For fine-tuning, we conducted 20 independent training runs on the same ImageNet-trained model, each with a distinct fixed seed. This approach ensured varied initializations for the new classifier head (adapted from 1000 output units to the 8 units needed for our classification task), for the dataloaders and different seed settings for essential libraries (e.g., *NumPy*, *PyTorch*, *fastai*, and CUDA). Each run comprised 3 cycles, each with 9 epochs, amounting to 27 epochs in total. The initial 6 epochs of each cycle focused on training the model’s classifier and the last convolutional layer before the classifier. This strategy was adopted to preserve low-level features learned from ImageNet, aligning with our goal to model human-like visual processing by retaining features from a dataset rich in natural visual variance. In the last 3 epochs of each cycle, the full model was trained, allowing for a comprehensive adaptation to our 8-way classification task. The learning rate for each cycle was dynamically determined using the learning rate finder algorithm from *fastai* at the start of each cycle. The `fit_one_cycle` method parameters were set with `lr_max` as the optimal learning rate multiplied by 10.

The fine-tuning dataset included a total of 32,000 images taken from the same datasets as these used to build the human dataset (see Stimuli), with an equal distribution of 4,000 images per category. Each category was divided to include half the images with their original background (split equally between colored and black and white) and half against a neutral gray background (also divided equally between colored and black and white). This approach allowed us to train the network to accurately categorize images irrespective of background variations and color cues.

This dual-phase training approach produced 20 uniquely fine-tuned versions of each model, ensuring robust adaptation to our classification task while preserving foundational ImageNet-acquired knowledge. The Adam optimizer with a Cross Entropy loss function was employed throughout both phases of training. Training was executed in parallel on 3 NVIDIA GeForce RTX 2080 GPUs. The models were implemented using *PyTorch* version 2.1.2 [Paszke et al., 2019], *TorchVision* version 0.16.2 [TorchVision maintainers and contributors, 2016], and *fastai* version 2.7.13 [Howard et al., 2018].

### Model Sizes and Connections

A comprehensive overview of the model architectures is provided in Fig. 2. Models that implemented at least one type of recurrent connection (i.e., top-down or within-layer recurrent mechanism) have an added time dimension due to the iterative nature of recurrent dynamic. Here, we set the total time-steps to 5 during both training and testing phases. A time-step here represents an end-to-end (from input to output) processing of the input image.

#### Top-Down Connections

Top-down mechanisms were implemented in three of the models used here: CT, CLT and BLT. The implementation of top-down mechanisms in the CT and CTL architectures, inspired by the CORnet model, and the BLT model, reveals distinct approaches to incorporating feedback connections in deep neural networks.

The BLT architecture incorporates top-down connections through transposed convolutional layers between consecutive layers, enabling the model to integrate abstract, higher-level information back into earlier stages of processing. The de-convolution and reintegration process could be seen as part of a predictive coding strategy, where the model minimizes prediction errors between higher level predictions and actual sensory input – akin to generative models that seek to enhance feature selectivity and reconstruct low-level sensory inputs from higher-level abstractions. The additive nature of the de-convolved signal in the implementation used here could lead to a direct influence of higher-level representations on lower-level processing, but offers less capacity for non-linear integration of the bottom-up and top-down signals compared to our other networks.

In CT and CLT models, on the other hand, each block is uniquely structured to integrate top-down inputs from all higher blocks, not just the adjacent one, through a specialized top-down pathway before being combined with the feedforward input through an additional convolutional layer. Such a design facilitates a more sophisticated non-linear integration of hierarchical information, allowing for the integration of bottom-up data with top-down information from all higher layers. This method fosters richer representation at each processing stage at the cost of increased computational complexity and a partial information loss due to channel averaging.

Specifically, in the CT and CLT models, a block processes a bottom-up input *x* (the output of the previous block at time-step *t −* 1) and combines it with top-down feedback *M*, derived from the aggregated outputs of higher-level blocks at time-step *t −* 1. In instances where top-down input is absent – such as during the initial time-step when no higher-level info is available – *M* is set to zero.

The feedforward path *FF* and the top-down path *TD* each involve convolution (*), Group Normalization (N), and ReLU activation (R):

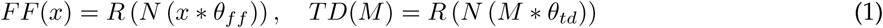

The weights *θ_ff_*and *θ_td_* represent distinct sets of learned weights for the feedforward and top-down convolutions, respectively.

The integrated top-down input *M* is formed by processing the outputs from higher layers *m*_1_*, m*_2_*, …, m_n_*:

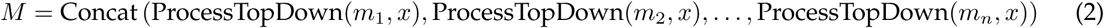

Given a top-down input tensor *m_i_* from layer *i* and a bottom-up input tensor *x*, the function ProcessLayer(*m_i_, x*) is defined as follows:

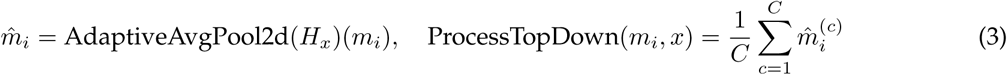

where *H_x_*is the height of the bottom-up input tensor *x*, *m*^ *_i_* is the tensor obtained after applying adaptive average pooling to *m_i_* with a target spatial dimension of *H_x_*, and the function computes the mean across the channel dimension of *m*^ *_i_*, resulting in a tensor with a single channel and same spatial dimensions as the bottom-up input.

The final output *y* of the block is generated by concatenating the feedforward and top-down outputs and applying convolution, normalization, and ReLU:

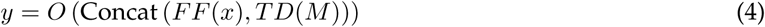

with *z* = Concat(*FF* (*x*)*, TD*(*M*)), where concatenation occurs along the channel dimension. The output operation *O* applies convolution, normalization, and ReLU activation on the concatenated tensor *z*:

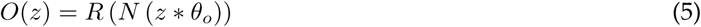

In this setup, the feedforward path (*FF*) processes input from the preceding block, while the top-down path (*TD*) integrates inputs from multiple higher layers, creating a comprehensive feedback mechanism. This design allows for a dynamic interplay of bottom-up perceptual data and high-level predictions. Top-down connections facilitate a backward flow of information, refining feedforward outputs with knowledge acquired from more abstract representations. This approach mirrors the concept of biologically plausible models, where visual information processing involves not only a feedforward pathway but also feedback loops that dynamically adjust and refine their understanding of visual inputs [van Bergen and Kriegeskorte, 2020].

#### Within-layer Connections

The CORblock RT block, used in the original implementation of CL [Kubilius et al., 2018]), was used here to model within-layer connections in models with within-layer connections only (i.e., CL, C V1-V1 and C IT-IT) to keep our custom implementations closer to the original. In these models, lateral connections are established within the block, where the output of a block at one timestep is used as an additional input in the next timestep (see [Kubilius et al., 2018] for more details). On the other hand, within-layer connections are implemented through temporal depth (i.e., through several iterations over the same block) in CORnet S, with skip connections that facilitates the preservation and integration of information across multiple passes of the same block, effectively approximating a very deep network architecture with shared weights. In contrast, lateral connections are implemented in BLT through specific convolutional layers that are dedicated to processing inputs from the same layer at previous timesteps. These layers are specifically tasked with processing the output of the same block from the previous timestep, effectively allowing the block to integrate its past state with the current input at the “cost” of additional parameters.

A different approach was used to model within-layer connections for C LT, due to the more complex shapes and different natures of the top-down, bottom-up and lateral inputs. For each CLT block, during the forward pass for an input *x* at a given timestep *t*, a previous state *s_t−_*_1_, and top-down input *td*, the output *y_t_* and the updated state *s_t_* can be described as follows:

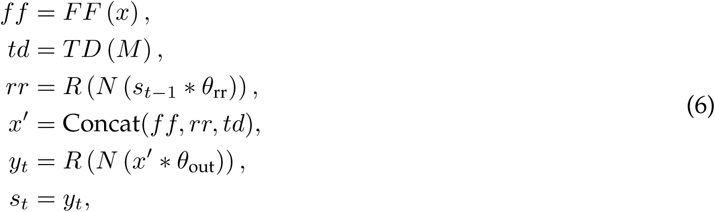

In this representation, *ff* corresponds to the output of the feedforward pass, computed by convolving the input *x* with the weights *θ*_ff_, followed by normalization *N* and ReLU activation *R*. The processed top-down input, *td*, is obtained by applying a sequence of operations defined by ProcessTopDown on the concatenated top-down input *M* (see sub-section *Top-Down Connections.* for more details). The term *rr* represents the within-layer processing of the previous state *s_t−_*_1_, convolved with *θ*_rr_ with kernel size and stride of 1 to preserve spatial dimensions, and then normalized and ReLU-activated. The concatenation operation Concat merges the outputs *ff*, *rr*, and *td* into a single tensor *x^i^* along the channel dimension. This concatenation preserves the spatial dimensions of the feature maps, and effectively pools the distinct features extracted from each individual path into a unified representation for subsequent layers. Finally, the output *y_t_* of the block at timestep *t* and the updated state *s_t_* are obtained by convolving *x^i^* with the output weights *θ*_out_, followed by normalization and ReLU activation. This output serves both as the response of the block at the current timestep and the state for the next timestep.

This recurrent structure enables the layer to integrate both current and previous activations, thereby enriching the artificial system with temporal dynamics and mimicking the recurrent mechanisms observed in biological visual systems ([Kietzmann et al., 2019, Kar et al., 2019]).

### Model Evaluation and Statistical Analysis

After training and fine-tuning, each model was tested against our test dataset. The test dataset included 800 black and white images (100 images for each of the same 8 categories) on a gray background. The dataset was presented to the network in a control or challenging condition (see *Challenging manipulations* for more details).

To evaluate the performance of each DNN model we employed a Top-1 Error Rate metric, which assesses classification accuracy based on the top prediction. Classification accuracy for each item in the test dataset was recorded for each model and each training seed, resulting in one confusion matrix per model across all the seeds.

## Results

In this study, we compared humans and DNN models on a categorisation task in order to investigate whether recurrent DNNs would better fit with human behaviour than feedforward DNNs. To capture these potential differences we developed a rich stimulus set that would be sensitive enough to allow for a fine-grained test of model fit to human data. The stimulus set quality was twofold. First, it implemented a variety of visual challenges (occlusion, phase scrambling, clutter), aimed at reproducing the diversity of natural scene complexity, while maintaining control over the degree of challenge. Second, it included a high number of object categories (eight categories: people, cats, birds, trees, bananas, buses, buildings and fire hydrants, see Fig 1), thereby requiring a focus upon multiple features and dimensions to solve the task. We chose to compare model types on their performance on a challenging recognition task, as recurrent processing in the brain has been extensively evidenced to play a role in the context of challenging visual inputs.

During the experiment, images were presented either manipulated (challenging conditions, 16 conditions) or non-manipulated (control condition, segmented objects on a plain background), for a total of 17 conditions to solve. Models and humans were asked to label the category the image belonged to on every trial, with human participants instructed to do so as fast and accurately as possible. We measured performance across visual manipulations, expecting that the advantages of adding various types of recurrent connections to our DNN models would especially show in the challenging conditions.

For humans only, half of the trials were masked. In masked trials, a phase scrambled noise pattern was presented shortly after the stimulus. This allowed for the selective impairing of recurrent processing, and served to confirm that the visual manipulations we implemented were linked to recurrence. With masked trials allowing for less contribution of recurrent processing in the brain, we expected that recurrent DNNs would show a better fit to non-masked trials, as compared to feedforward DNNs.

Different DNN models were trained and tested that covered a wide range of architectures and sizes (see Fig. 2). The architectures we focused on were either purely ***feedforward*** or contained ***recurrence***. Within our recurrent models, different types of recurrent connections were implemented. We used existing models from three families: *C*, *B*, and *VGG* ([Kubilius et al., 2018, Spoerer et al., 2017, Simonyan and Zisserman, 2015]). While the *VGG* family is purely feedforward and aims at maximising performance through depth, both the *C* and *B* families involve recurrence and are built as biologically plausible architectures. Conversely to the B models where all cases of feedback and lateral connectivity are explored, the C models included only a base, feedforward model (*C Z*, referred to hereafter as *C*), a fully laterally connected model (*C RT*, referred to hereafter as *CL*), and a high-performance, enhanced version of the latter (*C S*). To increase the spectrum of recurrent architectures in this family, we built ***four extra C architectures***: *C V1-V1*, *C IT-IT*, *CT* and *CLT*. On top of C and CL, these new models allowed us to separately look at the role of lateral and feedback connections in the C family too. In addition to architecture, we had large model size differences across our DNNs, with smaller models (e.g. C, 4.5m parameters) more than twenty times smaller than our bigger ones (e.g. VGG 16, 110m parameters). Overall, we used a range of models capable of dissociating between the effects of depth, size, and recurrent connectivities on performance. All our models underwent a similar regimen of training: first on ImageNet, then fine-tuned on a custom dataset of images from the eight selected categories.

### Visual manipulations trigger recurrent processing

To evaluate the effect of backward masking on task performance, two t-tests were ran to compare average accuracy and average reaction time (RT) with and without masking. Results showed a significantly lower average accuracy in the presence of masking (*t* = *−*3.12, *p* = 0.006, see Fig. 3A), but no significant difference in average RT (see Fig. 3C). Across manipulations, a wide range of accuracies were found, with some conditions linked to large increases in difficulty with masking, and others not. To elaborate on this difference, manipulations were split in two groups based on performance: an *easy* group (average accuracy *>* 0.9, 9 conditions) and a *hard* group (average accuracy *<* 0.9, 8 conditions), and compared average accuracy with and without masking within each group. As anticipated, this revealed significantly lower accuracy in the presence of masking in the hard manipulations (*t* = *−*4.47, *p* = 0.002) but not in the easy manipulations (*t* = *−*1.39, *p* = 0.2, see Fig. 3B).

**Figure 3:**
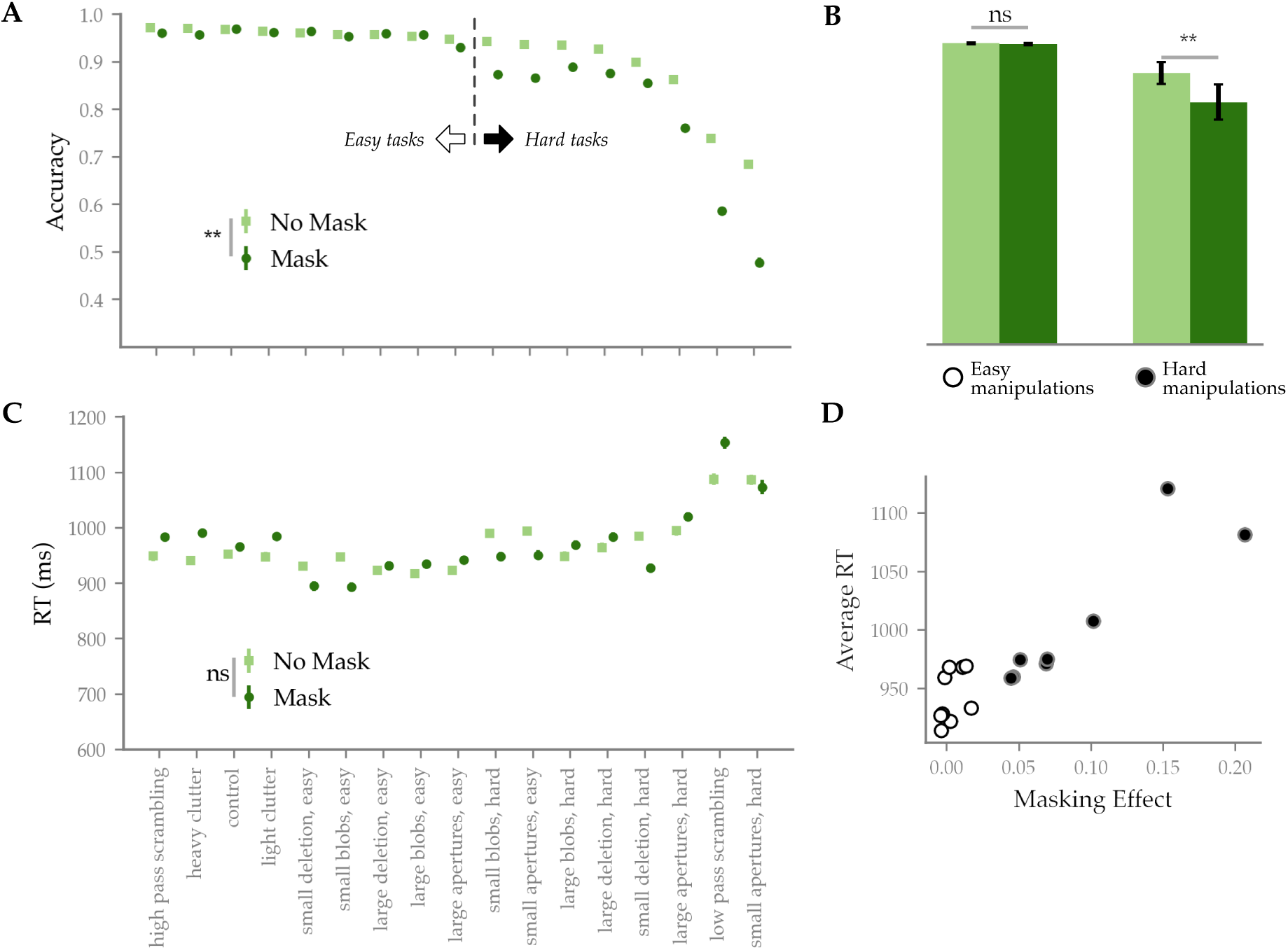
Human behavioural results. (A) Average accuracy across image manipulations, with and without masking. Manipulations are arranged along the x axis in order of average accuracy. A significant difference was found on average between the masked and non-masked condition. Manipulations were split into an easy and a hard group based on a 0.9 average accuracy threshold. (B) Average accuracy across masked and non masked trials, for the easy and hard manipulations. A significant masking effect was found only in the hard manipulations, confirming their interaction with recurrent processing. (C) Average RT across manipulations, with and without masking. The order on the x axis is the same as (A). No significant difference was found between average RT with and without masking. (D) Average RT and masking effect per condition, across manipulation difficulty groups. The two measures of recurrent processing are correlated (*r* = 0.93).

The masking effects found in our hard conditions indicate the implication of recurrent processing in solving the visual manipulations they contain. To support this interpretation, we turned to RT. Masking effect and RT are traditionally used to measure recurrence in visual recognition, but separately. Longer RTs are typically taken as indicators of extra, post-feedforward computations, while backward masking effects are direct evidence for the need of recurrent processing to solve the task. Here, we quantified the relationship of these two indicators by correlating the manipulation-wise average RT and backward masking effect (defined here as the average difference in accuracy between non masked and masked trials, see Fig. 3D).

We found a strong, positive correlation between average RT and masking effect (Pearson’s correlation *r* = 0.92, *p* = 7.04*e−*08), which remains strong even after the exclusion of the two most extremes performance values (*low-pass phase scrambling* and *many apertures, hard*, Pearson’s correlation *r* = 0.86, *p* = 4.5*e −* 05). The concurrence of both measures of recurrence in our results shows that our conditions covary on difficulty and need for recurrent processing.

### Model size drives average performance

Having established the role of recurrent processing in our stimulus set, we next asked how our models would perform on it. We started by comparing model average performances, in order to see whether added recurrence or larger model size would lead to an improvement at all. Average accuracy per model was collected for each of the 20 random initiation seeds (see Fig 4A). Performances were first compared across models by running a one-way ANOVA on accuracy with model as a main factor. Results showed a significant effect of models (*F* = 474.63, *p* = 5.64*e −* 163). Post-hoc comparisons of the models average accuracies showed significant differences across almost all pairs of models (Tukey test; all pair-wise comparisons significant, except: C - C V1-V1, C IT-IT - B, CT - BL, CLT - BT - BLT, CL - BL - BT, BT - BLT). In particular, VGG16 showed significantly higher accuracy than all models (*p <* 0.001, Tukey test). This reflects a striking difference in performance, with VGG16 on average 10% more accurate than other models.

**Figure 4:**
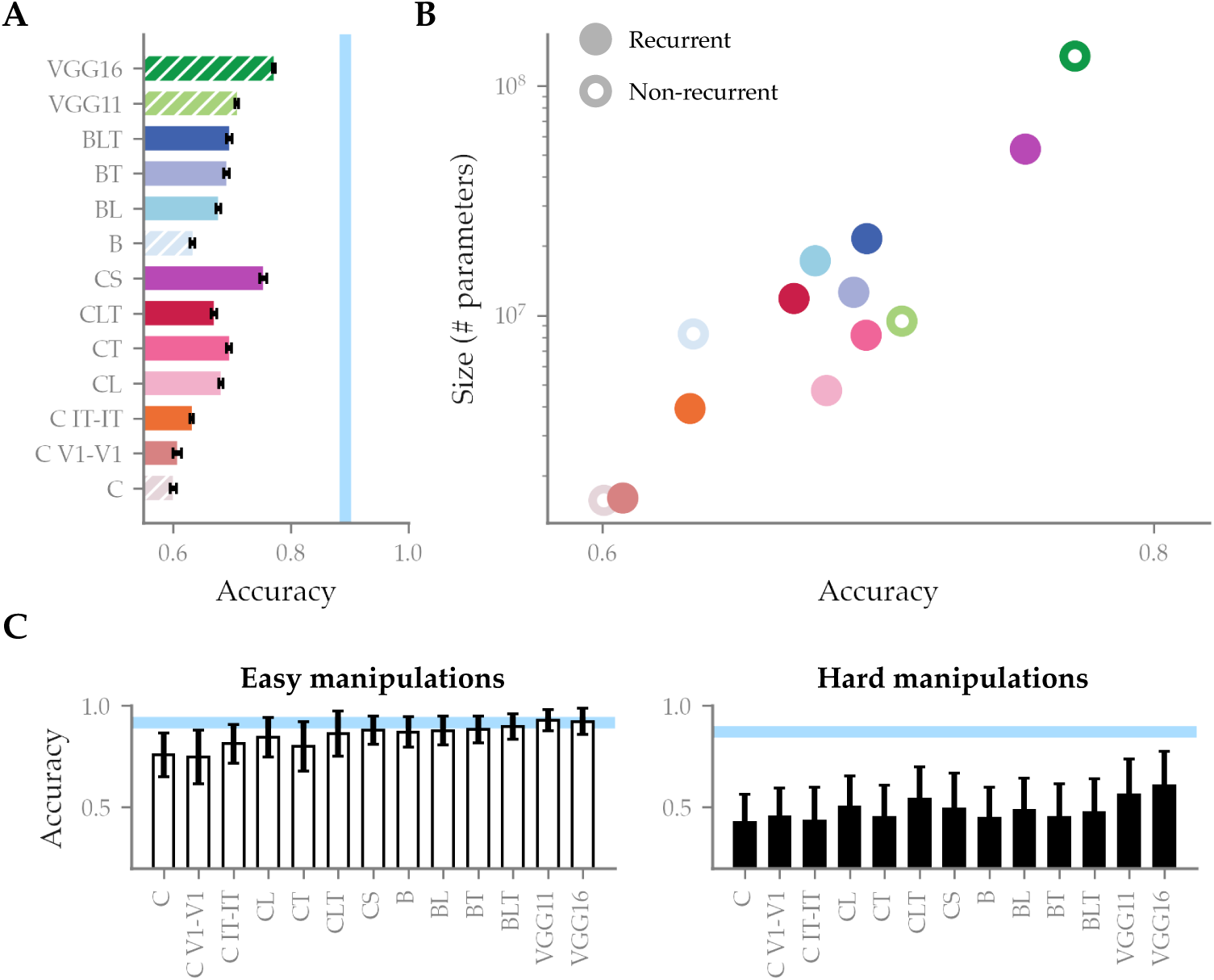
Average model performance. (A) Average model accuracy across all manipulations. Error bars show a 95% confidence interval around the mean calculated from the 20 random initialisations of each model. Hatched bars indicate non-recurrent models. The blue bar indicates average human performance on the task (0.89). (B) Average model performance per model size, in number of parameters (note the logarithmic scale of the y axis). Non recurrent models are indicated by a white-filled dot. (C) Average model performance on easy and hard manipulations separately. Blue bars indicate average human performance (easy manipulations: 0.92, hard manipulations: 0.87). Error bars show a confidence interval around the mean as in (A).

While the more recurrent DNNs (CL, CT, CLT, CS, BL, BT & BLT) seem to perform higher than their baseline, feedforward-only counterparts (CL, CT, CLT and CS *>* 95% CI of C; BT and BLT*>*= 95% CI of B), this improvement could also be explained by the increased size of these models. To check whether performance across models was driven by size, we correlated model parameter number with model average performance (see 4B). Results showed a large correlation (Pearson’s correlation *r* = 0.74, *p* = 0.003), pointing to **model size** as the main driver of the performances of our models.

With recurrence particularly helpful in challenging visual settings, we next checked whether this relationship with model size would hold across task difficulty. To this end, we ran a similar correlation, including only the easy or the hard manipulations, as defined above (see Fig. 3B). Results (see Fig. 4C) showed a stronger size-performance correlation in the hard conditions (Pearson’s correlation *r* = 0.68, *p* = 0.01) than in the easy conditions (Pearson’s correlation *r* = 0.51, *p* = 0.07).

### Better performing models are more consistent with human performance patterns across image manipulations

While performance is a good indicator of how well a given model can solve the complexity of our stimulus set, it does not take into account the variability in task difficulty across the challenging manipulations. A model could have a lower overall performance but a better condition-wise fit with humans. To look beyond accuracy, we asked which models most consistently fitted the pattern of performance of humans. We computed 17 average accuracy values (one for each manipulation) per model and for humans, and correlated the pattern of accuracy across manipulations between each model and the human pattern.

Results from this analysis (see Fig 5) show an overall high correlation (*>* 0.65), indicating that DNNs are on average consistent with humans and agree on which manipulation is easier or more difficult. Once again, models equipped with recurrent connectivity are more consistent than their feedforward counterpart: with the exception of *C V1-V1*, all recurrent models achieve larger correlation scores than their baseline equivalent (all recurrent *C* and *B* models with a mean correlation above the 95% CI of both baseline *C* and *B* models). Notably, VGG16 is the most consistent model (correlation of 0.85).

**Figure 5:**
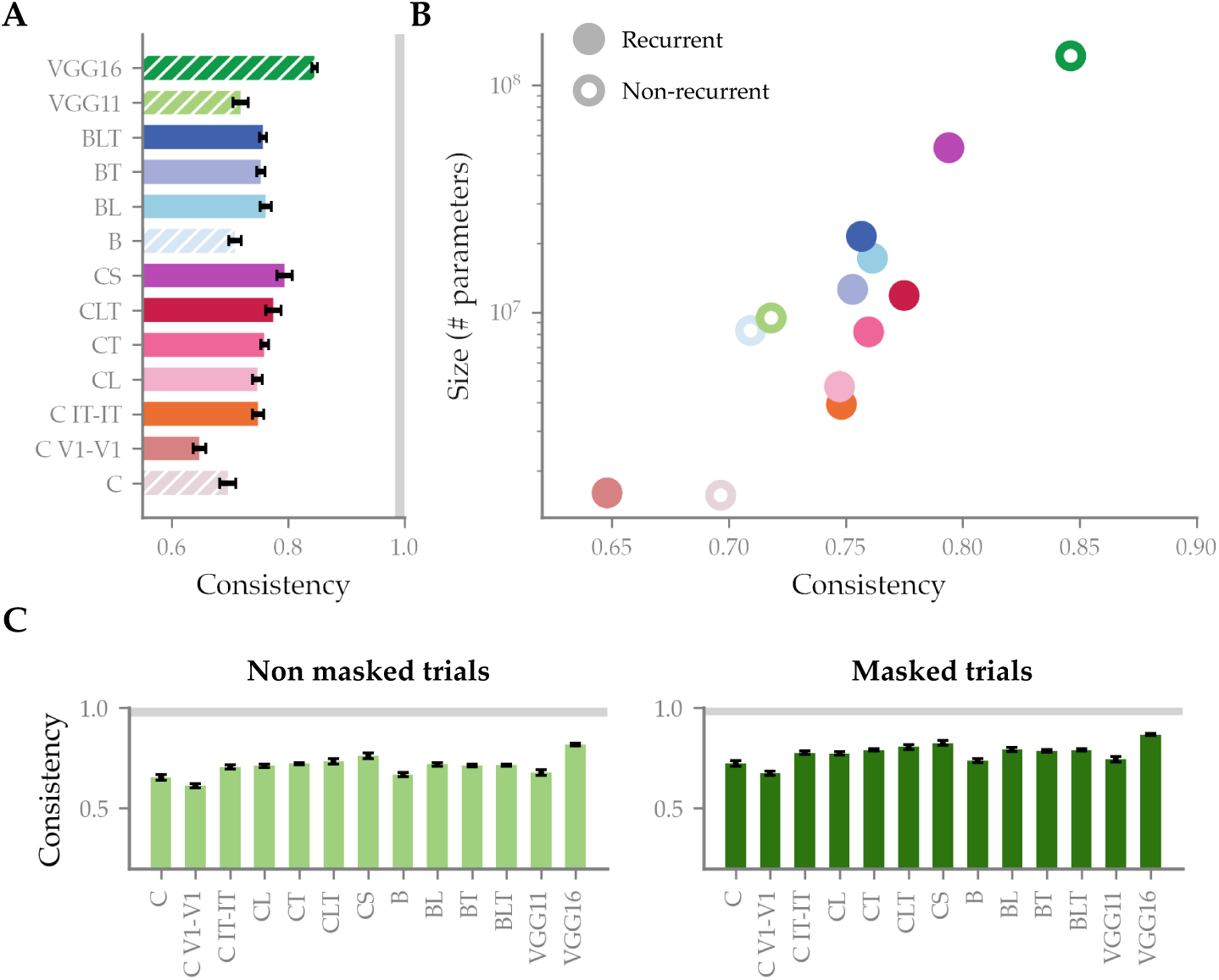
Consistency between model and human patterns of task performance. (A) Model consistencies, calculated as the correlations between human and model patterns of performance across manipulations. Error bars show a 95% confidence interval around the mean correlated calculated from the 20 random initialisations of each model. Hatched bars indicate non-recurrent models. The grey bar indicates the average split-half reliability of human accuracy across conditions (0.99) calculated over 100 iterations. (B) Model consistency with humans per model size, in number of parameters (note the logarithmic scale of the y axis). Non recurrent models are indicated by a white-filled dot. (C) Model consistencies on masked and non masked trials separately. Grey bars indicate average split-half reliability calculated as in (A) (reliability from non masked trials: 0.98, reliability from masked trials: 0.99). Error bars show a confidence interval around the mean as in (A).

While these results might show that, at approximately equivalent sizes, some recurrent models fare better on matching human behaviour than feedforward-only models (for instance, CT & CLT do better than VGG11 with approximately similar parameter numbers), model size could be the main driver of this effect. To test this, we correlated parameter number and consistency, and found a strong, positive relationship between them (Pearson’s correlation *r* = 0.75, *p* = 0.003), which again points to size as the main driver of model fit with human behaviour, above model architecture (see Fig. 5B).

Since it is expected that recurrent activity is impaired in the presence of backward masking, it can be predicted that recurrent models would be more consistent with humans in the absence of it, when recurrent activity unfolds normally. To test this, we computed similar correlations between size and consistency using trials with and trials without masking (see Fig. 5C). While an advantage of recurrent models in the absence of backward masking would have led to a decrease in the size-consistency relationship in non masked trials, we found similar correlations for masked trials (*r* = 0.72, *p* = 0.005) and unmasked trials (*r* = 0.79, *p* = 0.001). This indicates a strong influence of model size on consistency, regardless of the impairment or not of recurrent processing in humans.

### Recurrence types do not dissociate across visual manipulations

While size seems the main determinant of model consistency, manipulation-specific effects could remain: some manipulations could fall out of the human-DNN consensus, and instead be better solved by humans or by models. To check for this, we looked for outliers on the human-model agreement (see Fig. 6A). Two noticeable pairs of outliers were found. In the phase scrambling condition, the *low pass* and *high pass* pair, surprisingly, shows models performing better than expected on the former, and worse than expected on the latter. This clashes with the generally accepted notion that DNNs rely on high-frequency information to perform classification ([Geirhos et al., 2022, Avberšek et al., 2021]). In the occlusion condition, the *MS occluder high* and *MS blobs high* pair also falls out of distribution, with models performing surprisingly well on the former and surprisingly low on the latter. This might be due to the high number of black pixels in these manipulations (see Fig 1). The *occluder* images, indeed, have on average more black pixels than the *blobs* images (301e3 v. 137e3 black pixels per image, on average). These black pixels might create a visual transient that could distract participants and interfere with performance, which could explain the unexpectedly low human accuracy.

**Figure 6:**
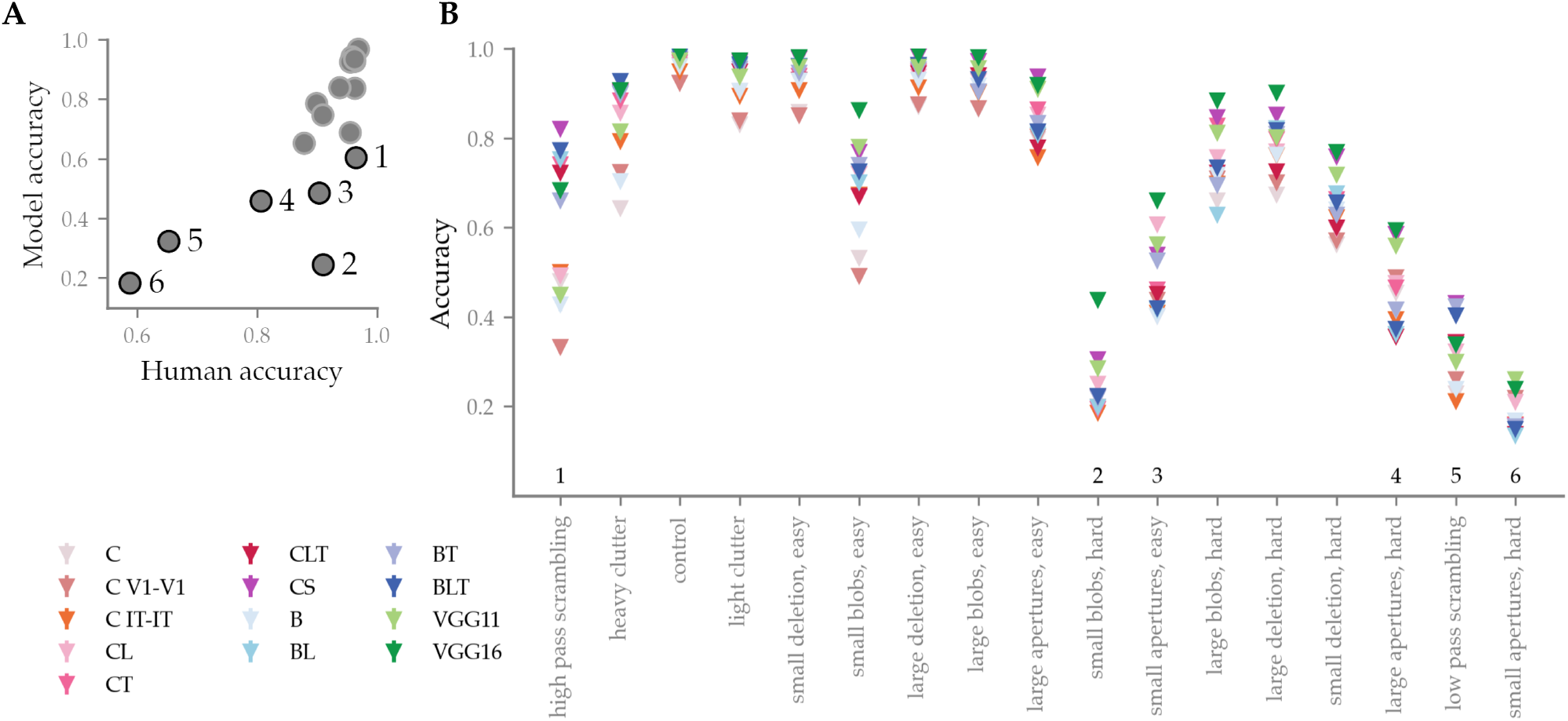
Performance across manipulations per models. (A) Model per human average accuracy across manipulations. Outliers are pointed out with reference to the numbers on (B). (B) Each diamond represents the average performance of a model on a given manipulation (bars on the diamonds indicate SEM). Manipulations are ordered on the x axis similarly to Fig 3A & C.

Better performing DNNs are on average more consistent with humans. This correspondence between general performance and consistency suggests that solving the categorisation task better leads to matching humans patterns better. However, this does not rule out model-specific effects, or connection-specific effects, whereby some conditions could be better solved by some models than others. There remains a lot of within-condition variability, and a given manipulation could be better solved by a model exhibiting, for instance, lateral recurrence, with that model not performing best overall (see Fig. 6B). To look for such effects and try to dissociate between types of recurrence, we checked whether the relationship between size and performance (see Fig 4) was repeated across conditions, and whether there were exceptions to size driving performance. For each condition, one vector of accuracy values was created with the average performance of each DNN model. Each vector (17 vectors in total, one per manipulation) was then correlated to the overall model order of performance.

Overall, the rankings of model performance across conditions seem to converge with the overall results: all conditions show a positive (Pearson’s *r >* 0.54, 0.76 on average) correlation with the order of model performance, with the only exception of *small apertures, hard* (correlation of 0.04). The latter deviates from the global trend, with baseline models reaching better accuracies than expected (e.g. VGG11 surpasses VGG16, B surpasses its B counterparts, C is the second best of the C models, see Fig. 6 for full details). While it is not yet clear why this manipulation falls out of the general trend, it is noticeably more difficult than average, and might show a *floor effect*. We further address this point in the discussion. Overall, the convergence of within-condition with overall model performance ranking indicates that models do not display strong condition-specific effects that could link architectural features to particular manipulation challenges.

### Depth, not recurrence, makes model mistakes more human-like

While model size seems to drive overall performance and consistency with human condition-wise behaviour, recurrence could make DNNs a better fit to human behaviour through a better match with the classification errors made by participants. We investigated this by comparing ***confusion matrices***. Confusion matrices are built by calculating the number of times each of the eight categories was given as a response, when each of the eight categories was presented. The result is a 8x8 matrix where each cell counts the number of responses to a given category. As a result, confusion matrices reflect the intrinsic overlap in feature representations that models and human participants use to perform classification. A confusion matrix was built for humans, and for each DNN model. The resulting matrices were correlated to quantify the agreement of each model with humans (matrices were correlated without diagonal values using Pearson’s R, see Fig 7).

**Figure 7:**
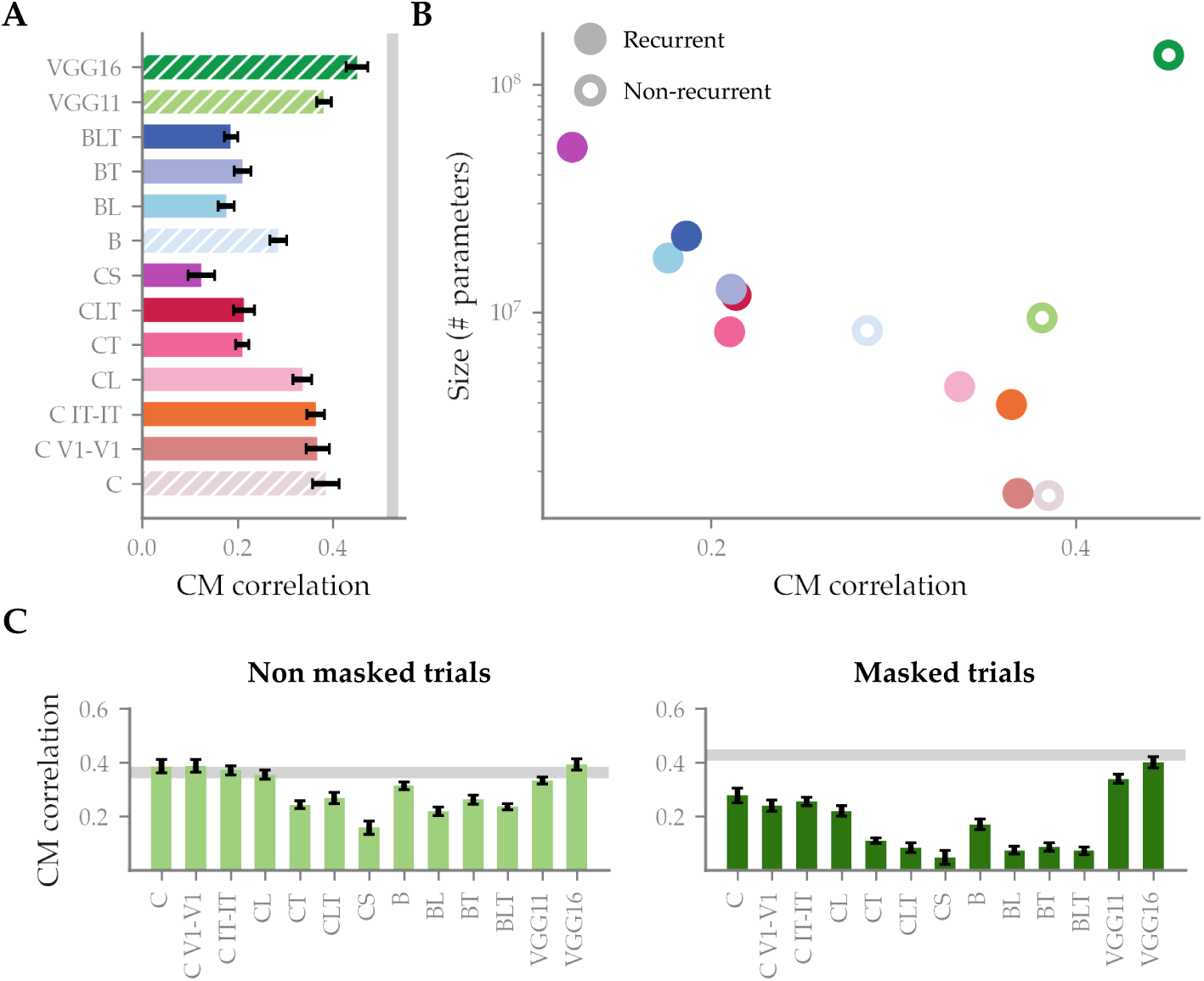
Correlation between confusion matrices from human and model behaviour. (A) Correlation with the average human confusion matrix, per model. Error bars show a 95% confidence interval around the mean confusion matrix correlation, calculated from the 20 random initialisations of each model. Hatched bars indicate non-recurrent models. The grey bar indicates the average split-half reliability of the human confusion matrix (0.52) calculated over 100 iterations. (B) Model correlation with human confusion matrix per model size, in number of parameters (note the logarithmic scale of the y axis). Non recurrent models are indicated by a white-filled dot. (C) Model correlation with human confusion matrix, extracted from masked and non masked trials separately. Grey bars indicate average split-half reliability calculated as in (A) (reliability from non masked trials: 0.36, reliability from masked trials: 0.43). Error bars show a confidence interval around the mean as in (A).

Strikingly, results show a disadvantage of adding recurrence, with models including more recurrent connections performing worse than their baseline counterparts in the two families of models that we included (see Fig. 7A-B). We confirmed these results by taking the more conservative approach of calculating correlations at the individual participant level and building confidence intervals around the mean of these correlations (see Fig. S4). Within recurrent models, the trend observed in previous analyses is reversed, with larger models performing worse than smaller ones.

To control for the effect of masking and the possibility that recurrent models would fit better with human behaviour in the absence of masking, we replicated this analysis on trials with and without masking (see Fig. 7C). Results indicate similar trends, with recurrent models showing a worse fit to human confusion matrices both with masking (Pearson’s correlation, *r* = 0.13 v. *r* = 0.3) and without masking (Pearson’s correlation, *r* = 0.28 v. *r* = 0.36). This is especially striking in comparison to the large, feedforward VGG16, which reaches noise ceiling-level correlations with both masked and non masked trials.

The results of our analyses show that model size drives performance and consistency with humans, with model architecture not playing a significant role. Although this fits with the idea that recurrent models are equivalent to time-unrolled feedforward models, we found that adding recurrence made models worse at replicating human classification errors. Additionally, we found that VGG 16, the deep, large, feedforward-only model, fits human data the best. Overall, we could not find any particular advantage of adding recurrent connectivity in DNNs as a better strategy to achieve a categorisation task in a more human-like fashion, as compared to depth and overall size.

## Discussion

We explored the role of recurrent processing in DNN models of visual recognition, testing the hypothesis that added recurrence in a model would make it solve object identity in a more human-like fashion. We designed a stimulus set with challenging visual features, and tested human participants with and without backward masking, to compare recurrent and non recurrent DNNs on their fit with human data in conditions where recurrence matters. Firstly, from the widespread observation that recurrence serves the visual system particularly in difficult visual conditions, we expected to find advantages of recurrent models in particularly challenging manipulations. Secondly, from the evidence that backward masking selectively impairs recurrent processing while leaving feedforward processing intact, we expected recurrent models to better fit with human behaviour in non-masked trials.

When looking at model accuracy, contrary to our expectations, we found a nuanced relationship between recurrence, model size, and performance. The inclusion of recurrent connections in DNNs improved performance, however, this improvement was tied to the overall size of the network. Larger models fared better on our manipulations than smaller ones, with VGG16 performing best. This pattern of results did not change for difficult conditions, where we expected recurrent DNNs to show an advantage.

In a comparable way, when looking at performance consistency across conditions, we did not find that recurrent models aligned better with human difficulty perception. Instead, a similar pattern emerged whereby larger models aligned more consistently with humans, with model size broadly predicting how well a given model would do. Additionally, the accuracy ranking of models collapsed when task difficulty reached a certain level (notably on the *MS blobs high occlusion* manipulation, see Recurrence types do not dissociate across visual manipulations), which seems to match with reports of DNNs failing to replicate human performance in highly difficult conditions ([Geirhos et al., 2018, Ghodrati et al., 2014]). Overall, we found an agreement between models and humans on task difficulty, in line with other reports showing a tendency of DNNs to generally fail when humans fail ([Kheradpisheh et al., 2016]). We also found this agreement to be performance-dependent, with larger models displaying more consistency than smaller ones, in line with other reports ([Lee and DiCarlo, 2023]). We did not find any difference between masked and non masked trials, while an advantage for recurrent models was expected for the latter.

We set up our experiment to be able to distinguish between different types of recurrence. By choosing and building DNNs that contained different types of recurrence in different layers, we aimed at finding manipulation-dependent effects of recurrent processing. However, we did not find any such distinctions in our results, as evidenced by the overall consensus of performance ranking across conditions (see Better performing models are more consistent with human performance patterns across image manipulations). This null result is surprising considering the ample evidence for region-specific perceptual phenomena in the brain. One could have expected, for instance, C V1-V1 to behave differently in conditions requiring figure-ground modulation (e.g. clutter tasks) given the known role of recurrent processing in V1 in this phenomenon ([Lamme et al., 1998, Self et al., 2013]). This result, combined with the general performance-dependent model consistency, shows that the recurrent connections we implemented do not critically change the behaviour of our models.

When considering task performance and consistency, our results align well with the notion that recurrent neural networks can be considered time-wrapped equivalents of size-matched feedforward networks ([van Bergen and Kriegeskorte, 2020, Liao and Poggio, 2020]). Recurrent architectures, though, could be considered superior because more brain-like. Furthermore, recurrence allows to increase the number of computations without an increase in the number of neurons. Thus, if we were to express model size in terms of the number of units rather than the number of parameters, then recurrence results in an improved performance without an increase in ‘size’.

However, even with this handy way out to promote recurrence as a solution with unique benefits, recurrence faces other challenges. Our examination of confusion matrices provided a finer-grained comparison of models, and showed, contrary to expectations, a drop in the alignment of recurrent models, specifically CS and BLT, with human behaviour. This corroborates reports showing that DNNs rely on different strategies to operate visual recognition ([Lonnqvist et al., 2019, Biscione and Bowers, 2023, Hosseini et al., 2017]). Importantly, we found a characteristic tradeoff between overall performance and fit with human data ([Fel et al., 2022]) within the recurrent model families C and B, but not VGG.

This challenges the observation that recurrent models better mimic the representational patterns of the brain than non-recurrent ones ([Nayebi et al., 2022, Kietzmann et al., 2019, Kar et al., 2019, Spoerer et al., 2020]). While recurrent models are reported to match better the brain’s functioning, our study suggests that they could simultaneously be worse models of human behaviour. Therefore, we emphasise behavioural fidelity as a metric for developing better models of human visual recognition.

Strikingly, although we used a design equipped for it, we found no clear dissociations between the effect of specific types of recurrence. Lateral connections did not induce specific effects compared to top-down connections, either in general or in a layer-dependent way. This lack of dissociation could mean that the implementation of recurrence within DNNs does not inherently alter the mechanisms of information processing. Rather than fundamentally reshaping how inputs are processed, recurrent processing, in its current implementation, seems to operate similarly to added depth, as a tool adding to the computational power of a network. This is in line with our observation that despite vast differences in architecture, the approach to visual recognition, as indexed by confusion matrices, remains relatively homogeneous for a given model size (see Fig S5).

It is noticeable that each of the recurrent connection in our models can be implemented in multiple ways. Here, we adhered to the common implementation as it is used by architectures such as C and B, yet, there is a wide range of other options and frequent developments in the field ([Nayebi et al., 2022, Wang and Hu, 2022, Linsley et al., 2019, Fukui et al., 2019, Lotter et al., 2017, Lazar et al., 2009, Konkle and Alvarez, 2023]). In addition, even though the visual challenges that we created were motivated by previous research that referred to a potential specific role of lateral versus top-down connections ([Rajaei et al., 2019, Seijdel et al., 2021, Bar et al., 2006]), there might be better manipulations to be found. Overall, before we can accept the hypothesis that the type of recurrence does not matter, further studies are needed with more refined, biologically plausible implementations of recurrence and with additional visual challenges.

The need to search further in the space of visual manipulations also finds support in the wide variety we found across our 17 manipulations. Interesting distinctions emerged, with seemingly similar manipulations bringing surprisingly different results. The implementation of many small occluders, for instance, brought two such experimental conditions: *MS occluder, high* and *MS blobs, high*. While in the former, both humans and models see significant drops in their performance, the latter shows disagreement, with humans finding it much less difficult. A similar distinction can be found by comparing phase scrambling in the *low pass* condition and *high pass* condition. Even in the absence of model-specific effects, our results highlight the behavioural variability that can be measured using a rich stimulus set.

In conclusion, contrary to our expectations, our findings highlight the limitations of recurrence as implemented in our models. Furthermore, they point out the overall discrepancy of visual processing in DNNs compared to humans. While we reproduce the general DNN ability to capture patterns of task difficulty, our results do not support that recurrent models outperform feedforward counterparts in capturing human visual recognition processes. As a consequence, we stress the importance of behavioural fidelity as a metric in developing new models, and emphasise the need for refined, biologically plausible implementations of recurrence.

## Supporting information

**Table 1: General model information**. Extra information about models parameter size, connections (lateral/feedback) and references.

**Table 2: Custom model extra information: C V1-V1**. Layer specific information (type, input and output shape, parameters, trainability) about **C V1-V1** (CORnet V1-V1).

**Table 3: Custom model extra information: C IT-IT**. Layer specific information (type, input and output shape, parameters, trainability) about **C IT-IT** (CORnet IT-IT).

**Table 4: Custom model extra information: CT**. Layer specific information (type, input and output shape, parameters, trainability) about **CT** (CORnet T).

**Table 5: Custom model extra information: CLT**. Layer-specific information (type, input and output shape, parameters, trainability) about **CLT** (CORnet LT).

**Table 6: Custom model extra information: VGG11**. Layer-specific information (type, input and output shape, parameters, trainability) about **VGG11**.

**Table 7: Average performance per task per model**. Detailed average performance per model and per task, as shown in figure 6.

**Figure S1: Stimulus manipulation details**. A schematic of how each of the challenging manipulation is implemented on the stimulus set.

**Figure S2: Colour-blind friendly performance pointplot**. A less colour-heavy reproduction of figure 6. The colour palette has been adapted for more contrast. Shape distinctions have been added as well.

**Figure S3: Model PCA plots**. PCA plots built from the category-wise confusion data of individual images, for each model and for human participants.

**Figure S4: Participant-level confusion matrix correlation**. Correlations between model and human confusion matrices, calculated at the participant level.

**Figure S5: Within-model confusion matrix correlations**. Correlation matrix (Pearson’s R) of the pair-wise confusion matrix correlations of all models.

## Notes

### Competing Interest Statement

The authors have declared no competing interest.

### Summary of Updates

Added masking analyses; added condition difficulty analyses; expanded all results figures; corrected noise ceiling

