## Supplementary materials for "Recurrent issues with deep neural network models of visual recognition"

### Supplementary material for: *Recurrent issues with deep neural networks of visual recognition*

#### 1 Models information

The following section contains extra information about the models included in the main paper. We provide some general information about all the models in table 1, followed by extra model-specific information for the six custom-made models (for other models from *CORnet*, *B* and *VGG*, see the respective original publications).

| Model name | Source | Size (parameters) | Feedforward | Lateral | Feedback |
|---|---|---|---|---|---|
| CORnet_Z | Kubilius et al. (2018) | 1,562,760 | Yes | No | No |
| CORnet_V1_V1 | Custom-made | 1,599,880 | Yes | L1-L1 | No |
| CORnet_IT_IT | Custom-made | 1,599,880 | Yes | L4-L4 | No |
| CORnet_RT | Kubilius et al. (2018) | 4,700,040 | Yes | Yes | No |
| CORnet_T | Custom-made | 11,480,904 | Yes | No | Yes |
| CORnet_LT | Custom-made | 11,480,904 | Yes | Yes | Yes |
| CORnet_S | Kubilius et al. (2018) | 52,907,720 | Yes | Yes | No |
| B_net | Spoerer et al. (2017) | 8,291,720 | Yes | No | No |
| BL_net | Spoerer et al. (2017) | 17,257,864 | Yes | Yes | No |
| BT_net | Spoerer et al. (2017) | 12,592,520 | Yes | No | Yes |
| BLT_net | Spoerer et al. (2017) | 21,558,664 | Yes | Yes | Yes |
| VGG11 | Simonyan and Zisserman (2015) | 9,426,696 | Yes | No | No |
| VGG16 | Simonyan and Zisserman (2015) | 134,301,768 | Yes | No | No |

Table 1: General information on the models used.

## C V1-V1

| Index | Name | Layer Type | Input Shape | Output Shape | Param # | Trainable |
|---|---|---|---|---|---|---|
| 2 | V1.conv_input | Conv2d | [1, 3, 224, 224] | [1, 64, 56, 56] | 9472 | True |
| 3 | V1.norm_input | GroupNorm | [1, 64, 56, 56] | [1, 64, 56, 56] | 128 | True |
| 4 | V1.nonlin_input | ReLU | [1, 64, 56, 56] | [1, 64, 56, 56] | - | - |
| 5 | V1.conv1 | Conv2d | [1, 64, 56, 56] | [1, 64, 56, 56] | 36864 | True |
| 6 | V1.norm1 | GroupNorm | [1, 64, 56, 56] | [1, 64, 56, 56] | 128 | True |
| 7 | V1.nonlin1 | ReLU | [1, 64, 56, 56] | [1, 64, 56, 56] | - | - |
| 8 | V1.output | Identity | [1, 64, 56, 56] | [1, 64, 56, 56] | - | - |
| 10 | V2.conv | Conv2d | [1, 64, 56, 56] | [1, 128, 56, 56] | 73856 | True |
| 11 | V2.nonlin | ReLU | [1, 128, 56, 56] | [1, 128, 56, 56] | - | - |
| 12 | V2.pool | MaxPool2d | [1, 128, 56, 56] | [1, 128, 28, 28] | - | - |
| 13 | V2.output | Identity | [1, 128, 28, 28] | [1, 128, 28, 28] | - | - |
| 15 | V4.conv | Conv2d | [1, 128, 28, 28] | [1, 256, 28, 28] | 295168 | True |
| 16 | V4.nonlin | ReLU | [1, 256, 28, 28] | [1, 256, 28, 28] | - | - |
| 17 | V4.pool | MaxPool2d | [1, 256, 28, 28] | [1, 256, 14, 14] | - | - |
| 18 | V4.output | Identity | [1, 256, 14, 14] | [1, 256, 14, 14] | - | - |
| 20 | IT.conv | Conv2d | [1, 256, 14, 14] | [1, 512, 14, 14] | 1180160 | True |
| 21 | IT.nonlin | ReLU | [1, 512, 14, 14] | [1, 512, 14, 14] | - | - |
| 22 | IT.pool | MaxPool2d | [1, 512, 14, 14] | [1, 512, 7, 7] | - | - |
| 23 | IT.output | Identity | [1, 512, 7, 7] | [1, 512, 7, 7] | - | - |
| 25 | decoder.avgpool | AdaptiveAvgPool2d | [1, 512, 7, 7] | [1, 512, 1, 1] | - | - |
| 26 | decoder.flatten | Flatten | [1, 512, 1, 1] | [1, 512] | - | - |
| 27 | decoder.linear | Linear | [1, 512] | [1, 8] | 4104 | True |
| Total params: 1599880 | | | | | | |
| Trainable params: 1599880 | | | | | | |
| Non-trainable params: 0 | | | | | | |

Table 2: Layer specific information about **C V1-V1** (CORnet V1-V1).

## C IT-IT

| Index | Name | Layer Type | Input Shape | Output Shape | Param # | Trainable |
|---|---|---|---|---|---|---|
| 2 | V1.conv | Conv2d | [1, 3, 224, 224] | [1, 64, 56, 56] | 9472 | True |
| 3 | V1.nonlin | ReLU | [1, 64, 56, 56] | [1, 64, 56, 56] | - | - |
| 4 | V1.pool | MaxPool2d | [1, 64, 56, 56] | [1, 64, 28, 28] | - | - |
| 5 | V1.output | Identity | [1, 64, 28, 28] | [1, 64, 28, 28] | - | - |
| 7 | V2.conv | Conv2d | [1, 64, 28, 28] | [1, 128, 28, 28] | 73856 | True |
| 8 | V2.nonlin | ReLU | [1, 128, 28, 28] | [1, 128, 28, 28] | - | - |
| 9 | V2.pool | MaxPool2d | [1, 128, 28, 28] | [1, 128, 14, 14] | - | - |
| 10 | V2.output | Identity | [1, 128, 14, 14] | [1, 128, 14, 14] | - | - |
| 12 | V4.conv | Conv2d | [1, 128, 14, 14] | [1, 256, 14, 14] | 295168 | True |
| 13 | V4.nonlin | ReLU | [1, 256, 14, 14] | [1, 256, 14, 14] | - | - |
| 14 | V4.pool | MaxPool2d | [1, 256, 14, 14] | [1, 256, 7, 7] | - | - |
| 15 | V4.output | Identity | [1, 256, 7, 7] | [1, 256, 7, 7] | - | - |
| 17 | IT.conv_input | Conv2d | [1, 256, 7, 7] | [1, 512, 7, 7] | 1180160 | True |
| 18 | IT.norm_input | GroupNorm | [1, 512, 7, 7] | [1, 512, 7, 7] | 1024 | True |
| 19 | IT.nonlin_input | ReLU | [1, 512, 7, 7] | [1, 512, 7, 7] | - | - |
| 20 | IT.conv1 | Conv2d | [1, 512, 7, 7] | [1, 512, 7, 7] | 2359296 | True |
| 21 | IT.norm1 | GroupNorm | [1, 512, 7, 7] | [1, 512, 7, 7] | 1024 | True |
| 22 | IT.nonlin1 | ReLU | [1, 512, 7, 7] | [1, 512, 7, 7] | - | - |
| 23 | IT.output | Identity | [1, 512, 7, 7] | [1, 512, 7, 7] | - | - |
| 25 | decoder.avgpool | AdaptiveAvgPool2d | [1, 512, 7, 7] | [1, 512, 1, 1] | - | - |
| 26 | decoder.flatten | Flatten | [1, 512, 1, 1] | [1, 512] | - | - |
| 27 | decoder.linear | Linear | [1, 512] | [1, 8] | 4104 | True |
| Total params: 3924104 | | | | | | |
| Trainable params: 3924104 | | | | | | |
| Non-trainable params: 0 | | | | | | |

Table 3: Layer specific information about **C IT-IT** (CORnet IT-IT).

## CT

| Index | Name | Layer Type | Input Shape | Output Shape | Param # | Trainable |
|---|---|---|---|---|---|---|
| 3 | V1.ff_pass.0 | Conv2d | [1, 3, 224, 224] | [1, 64, 56, 56] | 9472 | True |
| 4 | V1.ff_pass.1 | GroupNorm | [1, 64, 56, 56] | [1, 64, 56, 56] | 128 | True |
| 5 | V1.ff_pass.2 | ReLU | [1, 64, 56, 56] | [1, 64, 56, 56] | - | - |
| 7 | V1.td_pass.0 | Conv2d | [1, 3, 224, 224] | [1, 64, 56, 56] | 9472 | True |
| 8 | V1.td_pass.1 | GroupNorm | [1, 64, 56, 56] | [1, 64, 56, 56] | 128 | True |
| 9 | V1.td_pass.2 | ReLU | [1, 64, 56, 56] | [1, 64, 56, 56] | - | - |
| 11 | V1.out_pass.0 | Conv2d | [1, 128, 56, 56] | [1, 64, 56, 56] | 401472 | True |
| 12 | V1.out_pass.1 | GroupNorm | [1, 64, 56, 56] | [1, 64, 56, 56] | 128 | True |
| 13 | V1.out_pass.2 | ReLU | [1, 64, 56, 56] | [1, 64, 56, 56] | - | - |
| 14 | V1.output | Identity | [1, 64, 56, 56] | [1, 64, 56, 56] | - | - |
| 17 | V2.ff_pass.0 | Conv2d | [1, 64, 56, 56] | [1, 128, 28, 28] | 73856 | True |
| 18 | V2.ff_pass.1 | GroupNorm | [1, 128, 28, 28] | [1, 128, 28, 28] | 256 | True |
| 19 | V2.ff_pass.2 | ReLU | [1, 128, 28, 28] | [1, 128, 28, 28] | - | - |
| 21 | V2.td_pass.0 | Conv2d | [1, 2, 56, 56] | [1, 128, 28, 28] | 2432 | True |
| 22 | V2.td_pass.1 | GroupNorm | [1, 128, 28, 28] | [1, 128, 28, 28] | 256 | True |
| 23 | V2.td_pass.2 | ReLU | [1, 128, 28, 28] | [1, 128, 28, 28] | - | - |
| 25 | V2.out_pass.0 | Conv2d | [1, 256, 28, 28] | [1, 128, 28, 28] | 295040 | True |
| 26 | V2.out_pass.1 | GroupNorm | [1, 128, 28, 28] | [1, 128, 28, 28] | 256 | True |
| 27 | V2.out_pass.2 | ReLU | [1, 128, 28, 28] | [1, 128, 28, 28] | - | - |
| 28 | V2.output | Identity | [1, 128, 28, 28] | [1, 128, 28, 28] | - | - |
| 31 | V4.ff_pass.0 | Conv2d | [1, 128, 28, 28] | [1, 256, 14, 14] | 295168 | True |
| 32 | V4.ff_pass.1 | GroupNorm | [1, 256, 14, 14] | [1, 256, 14, 14] | 512 | True |
| 33 | V4.ff_pass.2 | ReLU | [1, 256, 14, 14] | [1, 256, 14, 14] | - | - |
| 35 | V4.td_pass.0 | Conv2d | [1, 1, 28, 28] | [1, 256, 14, 14] | 2560 | True |
| 36 | V4.td_pass.1 | GroupNorm | [1, 256, 14, 14] | [1, 256, 14, 14] | 512 | True |
| 37 | V4.td_pass.2 | ReLU | [1, 256, 14, 14] | [1, 256, 14, 14] | - | - |
| 39 | V4.out_pass.0 | Conv2d | [1, 512, 14, 14] | [1, 256, 14, 14] | 1179904 | True |
| 40 | V4.out_pass.1 | GroupNorm | [1, 256, 14, 14] | [1, 256, 14, 14] | 512 | True |
| 41 | V4.out_pass.2 | ReLU | [1, 256, 14, 14] | [1, 256, 14, 14] | - | - |
| 42 | V4.output | Identity | [1, 256, 14, 14] | [1, 256, 14, 14] | - | - |
| 45 | IT.ff_pass.0 | Conv2d | [1, 256, 14, 14] | [1, 512, 7, 7] | 1180160 | True |
| 46 | IT.ff_pass.1 | GroupNorm | [1, 512, 7, 7] | [1, 512, 7, 7] | 1024 | True |
| 47 | IT.ff_pass.2 | ReLU | [1, 512, 7, 7] | [1, 512, 7, 7] | - | - |
| 49 | IT.td_pass.0 | Conv2d | [1, 1, 14, 14] | [1, 512, 7, 7] | 5120 | True |
| 50 | IT.td_pass.1 | GroupNorm | [1, 512, 7, 7] | [1, 512, 7, 7] | 1024 | True |
| 51 | IT.td_pass.2 | ReLU | [1, 512, 7, 7] | [1, 512, 7, 7] | - | - |
| 53 | IT.out_pass.0 | Conv2d | [1, 1024, 7, 7] | [1, 512, 7, 7] | 4719104 | True |
| 54 | IT.out_pass.1 | GroupNorm | [1, 512, 7, 7] | [1, 512, 7, 7] | 1024 | True |
| 55 | IT.out_pass.2 | ReLU | [1, 512, 7, 7] | [1, 512, 7, 7] | - | - |
| 56 | IT.output | Identity | [1, 512, 7, 7] | [1, 512, 7, 7] | - | - |
| 58 | decoder.avgpool | AdaptiveAvgPool2d | [1, 512, 7, 7] | [1, 512, 1, 1] | - | - |
| 59 | decoder.flatten | Flatten | [1, 512, 1, 1] | [1, 512] | - | - |
| 60 | decoder.linear | Linear | [1, 512] | [1, 8] | 4104 | True |
| Total params: 8183624 | | | | | | |
| Trainable params: 8183624 | | | | | | |
| Non-trainable params: 0 | | | | | | |

Table 4: Layer specific information about **CT**.

#### CLT

| Index | Name | Layer Type | Input Shape | Output Shape | Param # | Trainable |
|---|---|---|---|---|---|---|
| 3 | V1.ff_pass.0 | Conv2d | [1, 3, 224, 224] | [1, 64, 56, 56] | 9472 | True |
| 4 | V1.ff_pass.1 | GroupNorm | [1, 64, 56, 56] | [1, 64, 56, 56] | 128 | True |
| 5 | V1.ff_pass.2 | ReLU | [1, 64, 56, 56] | [1, 64, 56, 56] | - | - |
| 7 | V1.rr_pass.0 | Conv2d | [1, 64, 56, 56] | [1, 64, 56, 56] | 4160 | True |
| 8 | V1.rr_pass.1 | GroupNorm | [1, 64, 56, 56] | [1, 64, 56, 56] | 128 | True |
| 9 | V1.rr_pass.2 | ReLU | [1, 64, 56, 56] | [1, 64, 56, 56] | - | - |
| 10 | V1.rr_pass.3 | Dropout | [1, 64, 56, 56] | [1, 64, 56, 56] | - | - |
| 12 | V1.td_pass.0 | Conv2d | [1, 3, 224, 224] | [1, 64, 56, 56] | 9472 | True |
| 13 | V1.td_pass.1 | GroupNorm | [1, 64, 56, 56] | [1, 64, 56, 56] | 128 | True |
| 14 | V1.td_pass.2 | ReLU | [1, 64, 56, 56] | [1, 64, 56, 56] | - | - |
| 16 | V1.out_pass.0 | Conv2d | [1, 192, 56, 56] | [1, 64, 56, 56] | 602176 | True |
| 17 | V1.out_pass.1 | GroupNorm | [1, 64, 56, 56] | [1, 64, 56, 56] | 128 | True |
| 18 | V1.out_pass.2 | ReLU | [1, 64, 56, 56] | [1, 64, 56, 56] | - | - |
| 19 | V1.output | Identity | [1, 64, 56, 56] | [1, 64, 56, 56] | - | - |
| 22 | V2.ff_pass.0 | Conv2d | [1, 64, 56, 56] | [1, 128, 28, 28] | 73856 | True |
| 23 | V2.ff_pass.1 | GroupNorm | [1, 128, 28, 28] | [1, 128, 28, 28] | 256 | True |
| 24 | V2.ff_pass.2 | ReLU | [1, 128, 28, 28] | [1, 128, 28, 28] | - | - |
| 26 | V2.rr_pass.0 | Conv2d | [1, 128, 28, 28] | [1, 128, 28, 28] | 16512 | True |
| 27 | V2.rr_pass.1 | GroupNorm | [1, 128, 28, 28] | [1, 128, 28, 28] | 256 | True |
| 28 | V2.rr_pass.2 | ReLU | [1, 128, 28, 28] | [1, 128, 28, 28] | - | - |
| 29 | V2.rr_pass.3 | Dropout | [1, 128, 28, 28] | [1, 128, 28, 28] | - | - |
| 31 | V2.td_pass.0 | Conv2d | [1, 2, 56, 56] | [1, 128, 28, 28] | 2432 | True |
| 32 | V2.td_pass.1 | GroupNorm | [1, 128, 28, 28] | [1, 128, 28, 28] | 256 | True |
| 33 | V2.td_pass.2 | ReLU | [1, 128, 28, 28] | [1, 128, 28, 28] | - | - |
| 35 | V2.out_pass.0 | Conv2d | [1, 384, 28, 28] | [1, 128, 28, 28] | 442496 | True |
| 36 | V2.out_pass.1 | GroupNorm | [1, 128, 28, 28] | [1, 128, 28, 28] | 256 | True |
| 37 | V2.out_pass.2 | ReLU | [1, 128, 28, 28] | [1, 128, 28, 28] | - | - |
| 38 | V2.output | Identity | [1, 128, 28, 28] | [1, 128, 28, 28] | - | - |
| 41 | V4.ff_pass.0 | Conv2d | [1, 128, 28, 28] | [1, 256, 14, 14] | 295168 | True |
| 42 | V4.ff_pass.1 | GroupNorm | [1, 256, 14, 14] | [1, 256, 14, 14] | 512 | True |
| 43 | V4.ff_pass.2 | ReLU | [1, 256, 14, 14] | [1, 256, 14, 14] | - | - |
| 45 | V4.rr_pass.0 | Conv2d | [1, 256, 14, 14] | [1, 256, 14, 14] | 65792 | True |
| 46 | V4.rr_pass.1 | GroupNorm | [1, 256, 14, 14] | [1, 256, 14, 14] | 512 | True |
| 47 | V4.rr_pass.2 | ReLU | [1, 256, 14, 14] | [1, 256, 14, 14] | - | - |
| 48 | V4.rr_pass.3 | Dropout | [1, 256, 14, 14] | [1, 256, 14, 14] | - | - |
| 50 | V4.td_pass.0 | Conv2d | [1, 1, 28, 28] | [1, 256, 14, 14] | 2560 | True |
| 51 | V4.td_pass.1 | GroupNorm | [1, 256, 14, 14] | [1, 256, 14, 14] | 512 | True |
| 52 | V4.td_pass.2 | ReLU | [1, 256, 14, 14] | [1, 256, 14, 14] | - | - |
| 54 | V4.out_pass.0 | Conv2d | [1, 768, 14, 14] | [1, 256, 14, 14] | 1769728 | True |
| 55 | V4.out_pass.1 | GroupNorm | [1, 256, 14, 14] | [1, 256, 14, 14] | 512 | True |
| 56 | V4.out_pass.2 | ReLU | [1, 256, 14, 14] | [1, 256, 14, 14] | - | - |
| 57 | V4.output | Identity | [1, 256, 14, 14] | [1, 256, 14, 14] | - | - |
| 60 | IT.ff_pass.0 | Conv2d | [1, 256, 14, 14] | [1, 512, 7, 7] | 1180160 | True |
| 61 | IT.ff_pass.1 | GroupNorm | [1, 512, 7, 7] | [1, 512, 7, 7] | 1024 | True |
| 62 | IT.ff_pass.2 | ReLU | [1, 512, 7, 7] | [1, 512, 7, 7] | - | - |
| 64 | IT.rr_pass.0 | Conv2d | [1, 512, 7, 7] | [1, 512, 7, 7] | 262656 | True |
| 65 | IT.rr_pass.1 | GroupNorm | [1, 512, 7, 7] | [1, 512, 7, 7] | 1024 | True |
| 66 | IT.rr_pass.2 | ReLU | [1, 512, 7, 7] | [1, 512, 7, 7] | - | - |
| 67 | IT.rr_pass.3 | Dropout | [1, 512, 7, 7] | [1, 512, 7, 7] | - | - |
| 69 | IT.td_pass.0 | Conv2d | [1, 1, 14, 14] | [1, 512, 7, 7] | 5120 | True |
| 70 | IT.td_pass.1 | GroupNorm | [1, 512, 7, 7] | [1, 512, 7, 7] | 1024 | True |
| 71 | IT.td_pass.2 | ReLU | [1, 512, 7, 7] | [1, 512, 7, 7] | - | - |
| 73 | IT.out_pass.0 | Conv2d | [1, 1536, 7, 7] | [1, 512, 7, 7] | 7078400 | True |
| 74 | IT.out_pass.1 | GroupNorm | [1, 512, 7, 7] | [1, 512, 7, 7] | 1024 | True |
| 75 | IT.out_pass.2 | ReLU | [1, 512, 7, 7] | [1, 512, 7, 7] | - | - |
| 76 | IT.output | Identity | [1, 512, 7, 7] | [1, 512, 7, 7] | - | - |
| 78 | decoder.avgpool | AdaptiveAvgPool2d | [1, 512, 7, 7] | [1, 512, 1, 1] | - | - |
| 79 | decoder.flatten | Flatten | [1, 512, 1, 1] | [1, 512] | - | - |
| 80 | decoder.linear | Linear | [1, 512] | [1, 8] | 4104 | True |
| Total params: 11831944 | | | | | | |
| Trainable params: 11831944 | | | | | | |
| Non-trainable params: 0 | | | | | | |

Table 5: Layer-specific information about **CLT**.

#### VGG11

| Index | Name | Layer Type | Input Shape | Output Shape | Param # | Trainable |
|---|---|---|---|---|---|---|
| 2 | features.0 | Conv2d | [1, 3, 224, 224] | [1, 64, 224, 224] | 1792 | True |
| 3 | features.1 | BatchNorm2d | [1, 64, 224, 224] | [1, 64, 224, 224] | 128 | True |
| 4 | features.2 | ReLU | [1, 64, 224, 224] | [1, 64, 224, 224] | - | - |
| 5 | features.3 | MaxPool2d | [1, 64, 224, 224] | [1, 64, 112, 112] | - | - |
| 6 | features.4 | Conv2d | [1, 64, 112, 112] | [1, 128, 112, 112] | 73856 | True |
| 7 | features.5 | BatchNorm2d | [1, 128, 112, 112] | [1, 128, 112, 112] | 256 | True |
| 8 | features.6 | ReLU | [1, 128, 112, 112] | [1, 128, 112, 112] | - | - |
| 9 | features.7 | MaxPool2d | [1, 128, 112, 112] | [1, 128, 56, 56] | - | - |
| 10 | features.8 | Conv2d | [1, 128, 56, 56] | [1, 256, 56, 56] | 295168 | True |
| 11 | features.9 | BatchNorm2d | [1, 256, 56, 56] | [1, 256, 56, 56] | 512 | True |
| 12 | features.10 | ReLU | [1, 256, 56, 56] | [1, 256, 56, 56] | - | - |
| 13 | features.11 | Conv2d | [1, 256, 56, 56] | [1, 256, 56, 56] | 590080 | True |
| 14 | features.12 | BatchNorm2d | [1, 256, 56, 56] | [1, 256, 56, 56] | 512 | True |
| 15 | features.13 | ReLU | [1, 256, 56, 56] | [1, 256, 56, 56] | - | - |
| 16 | features.14 | MaxPool2d | [1, 256, 56, 56] | [1, 256, 28, 28] | - | - |
| 17 | features.15 | Conv2d | [1, 256, 28, 28] | [1, 512, 28, 28] | 1180160 | True |
| 18 | features.16 | BatchNorm2d | [1, 512, 28, 28] | [1, 512, 28, 28] | 1024 | True |
| 19 | features.17 | ReLU | [1, 512, 28, 28] | [1, 512, 28, 28] | - | - |
| 20 | features.18 | Conv2d | [1, 512, 28, 28] | [1, 512, 28, 28] | 2359808 | True |
| 21 | features.19 | BatchNorm2d | [1, 512, 28, 28] | [1, 512, 28, 28] | 1024 | True |
| 22 | features.20 | ReLU | [1, 512, 28, 28] | [1, 512, 28, 28] | - | - |
| 23 | features.21 | MaxPool2d | [1, 512, 28, 28] | [1, 512, 14, 14] | - | - |
| 24 | features.22 | Conv2d | [1, 512, 14, 14] | [1, 512, 14, 14] | 2359808 | True |
| 25 | features.23 | BatchNorm2d | [1, 512, 14, 14] | [1, 512, 14, 14] | 1024 | True |
| 26 | features.24 | ReLU | [1, 512, 14, 14] | [1, 512, 14, 14] | - | - |
| 27 | features.25 | Conv2d | [1, 512, 14, 14] | [1, 512, 14, 14] | 2359808 | True |
| 28 | features.26 | BatchNorm2d | [1, 512, 14, 14] | [1, 512, 14, 14] | 1024 | True |
| 29 | features.27 | ReLU | [1, 512, 14, 14] | [1, 512, 14, 14] | - | - |
| 30 | features.28 | MaxPool2d | [1, 512, 14, 14] | [1, 512, 7, 7] | - | - |
| 31 | classifier | Linear | [1, 25088] | [1, 8] | 200712 | True |
| Total params: 9426696 | | | | | | |
| Trainable params: 9426696 | | | | | | |
| Non-trainable params: 0 | | | | | | |

Table 6: Layer-specific information about **VGG11**.

### 2 Model performance table

Table 7 below summarises the data used to make figure 6 of the article. It contains the average accuracy per task, for all models as well as for humans.

| | Control | Light clutter | Heavy clutter | High pass | Low pass | Large deletion low | Large deletion high | Large apertures low | Large apertures high |
|---|---|---|---|---|---|---|---|---|---|
| CORnet_Z | 0.93 | 0.842 | 0.628 | 0.53 | 0.204 | 0.873 | 0.678 | 0.807 | 0.45 |
| CORnet_V1_V1 | 0.905 | 0.841 | 0.759 | 0.291 | 0.257 | 0.886 | 0.695 | 0.798 | 0.481 |
| CORnet_IT_IT | 0.947 | 0.898 | 0.783 | 0.497 | 0.21 | 0.92 | 0.767 | 0.749 | 0.383 |
| CORnet_RT | 0.967 | 0.91 | 0.858 | 0.485 | 0.326 | 0.94 | 0.778 | 0.847 | 0.466 |
| CORnet_T | 0.97 | 0.936 | 0.896 | 0.517 | 0.35 | 0.941 | 0.765 | 0.788 | 0.379 |
| CORnet_LT | 0.984 | 0.949 | 0.909 | 0.692 | 0.337 | 0.951 | 0.776 | 0.856 | 0.413 |
| CORnet_S | 0.993 | 0.972 | 0.903 | 0.821 | 0.432 | 0.983 | 0.852 | 0.938 | 0.585 |
| B_net | 0.963 | 0.906 | 0.686 | 0.362 | 0.237 | 0.932 | 0.77 | 0.816 | 0.41 |
| BL_net | 0.98 | 0.958 | 0.905 | 0.751 | 0.343 | 0.958 | 0.819 | 0.814 | 0.372 |
| BT_net | 0.976 | 0.954 | 0.897 | 0.627 | 0.405 | 0.955 | 0.81 | 0.839 | 0.444 |
| BLT_net | 0.981 | 0.962 | 0.916 | 0.78 | 0.399 | 0.961 | 0.833 | 0.813 | 0.37 |
| VGG11 | 0.974 | 0.937 | 0.805 | 0.414 | 0.275 | 0.956 | 0.801 | 0.903 | 0.546 |
| VGG16 | 0.99 | 0.97 | 0.904 | 0.673 | 0.316 | 0.985 | 0.906 | 0.914 | 0.596 |
| Humans | 0.969 | 0.963 | 0.963 | 0.965 | 0.653 | 0.958 | 0.899 | 0.938 | 0.807 |

| | Large blobs high | Large blobs low | Small deletion low | Small deletion high | Small blobs low | Small blobs high | Small apertures low | Small apertures high |
|---|---|---|---|---|---|---|---|---|
| CORnet_Z | 0.664 | 0.872 | 0.862 | 0.573 | 0.518 | 0.183 | 0.373 | 0.209 |
| CORnet_V1_V1 | 0.707 | 0.86 | 0.848 | 0.618 | 0.472 | 0.218 | 0.415 | 0.221 |
| CORnet_IT_IT | 0.696 | 0.905 | 0.898 | 0.644 | 0.652 | 0.191 | 0.41 | 0.159 |
| CORnet_RT | 0.758 | 0.928 | 0.929 | 0.609 | 0.756 | 0.253 | 0.591 | 0.207 |
| CORnet_T | 0.692 | 0.93 | 0.941 | 0.662 | 0.832 | 0.283 | 0.592 | 0.164 |
| CORnet_LT | 0.722 | 0.929 | 0.951 | 0.634 | 0.764 | 0.204 | 0.539 | 0.187 |
| CORnet_S | 0.847 | 0.972 | 0.979 | 0.76 | 0.769 | 0.305 | 0.54 | 0.156 |
| B_net | 0.746 | 0.916 | 0.928 | 0.637 | 0.58 | 0.226 | 0.432 | 0.171 |
| BL_net | 0.616 | 0.898 | 0.964 | 0.675 | 0.661 | 0.199 | 0.405 | 0.135 |
| BT_net | 0.709 | 0.904 | 0.948 | 0.652 | 0.754 | 0.209 | 0.527 | 0.168 |
| BLT_net | 0.716 | 0.942 | 0.957 | 0.66 | 0.674 | 0.234 | 0.423 | 0.15 |
| VGG11 | 0.808 | 0.954 | 0.963 | 0.701 | 0.784 | 0.308 | 0.552 | 0.256 |
| VGG16 | 0.88 | 0.983 | 0.982 | 0.777 | 0.865 | 0.425 | 0.661 | 0.261 |
| Humans | 0.91 | 0.956 | 0.963 | 0.879 | 0.955 | 0.91 | 0.904 | 0.588 |

Table 7: Average performance per task per model.

### 3 Supplementary figures

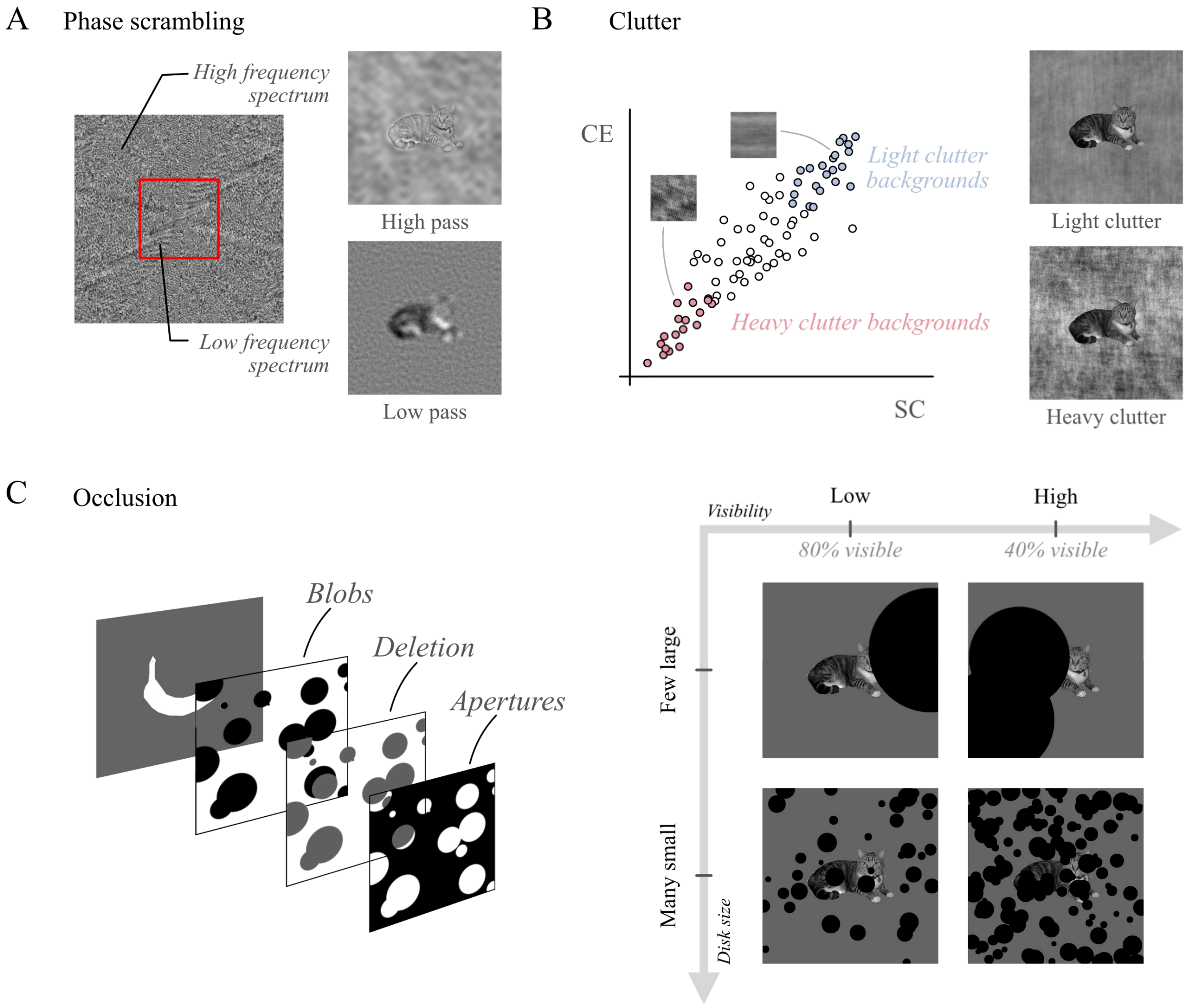


Figure S1: **Stimulus manipulations**. The stimulus set included eight object categories of 10 objects each, passed through a total of 16 visual manipulations. Objects in the images were rendered challenging by applying either *phase scrambling*, *clutter*, or *occlusion*. (A) Phase scrambling was applied by replacing the phase spectrum of images with random noise, on either sides of a 1.5 cpd threshold (red square not actual size). (B) Clutter was created following the method described in [**?**], whereby phase-scrambled versions of natural scenes taken from the MS COCO dataset were ranked on their *spatial coherence* and *contrast energy* values. Subsets of the most and less cluttered of a large number of such images were taken as light- and heavy-cluttered backgrounds for the stimuli. (C) Images were lightly or heavily occluded (percentage of object left visible: 80% and 40%, respectively), using many small or a few large disks. This was done in one of three possible fashions: adding black blobs, adding blobs the colour of the background (deletion) adding a full black occluder with disk-like apertures.

##### 3.1 Colour-blind friendly performance point plot

Supplementary figure S2 reproduces figure 6 in a less colour-heavy fashion.

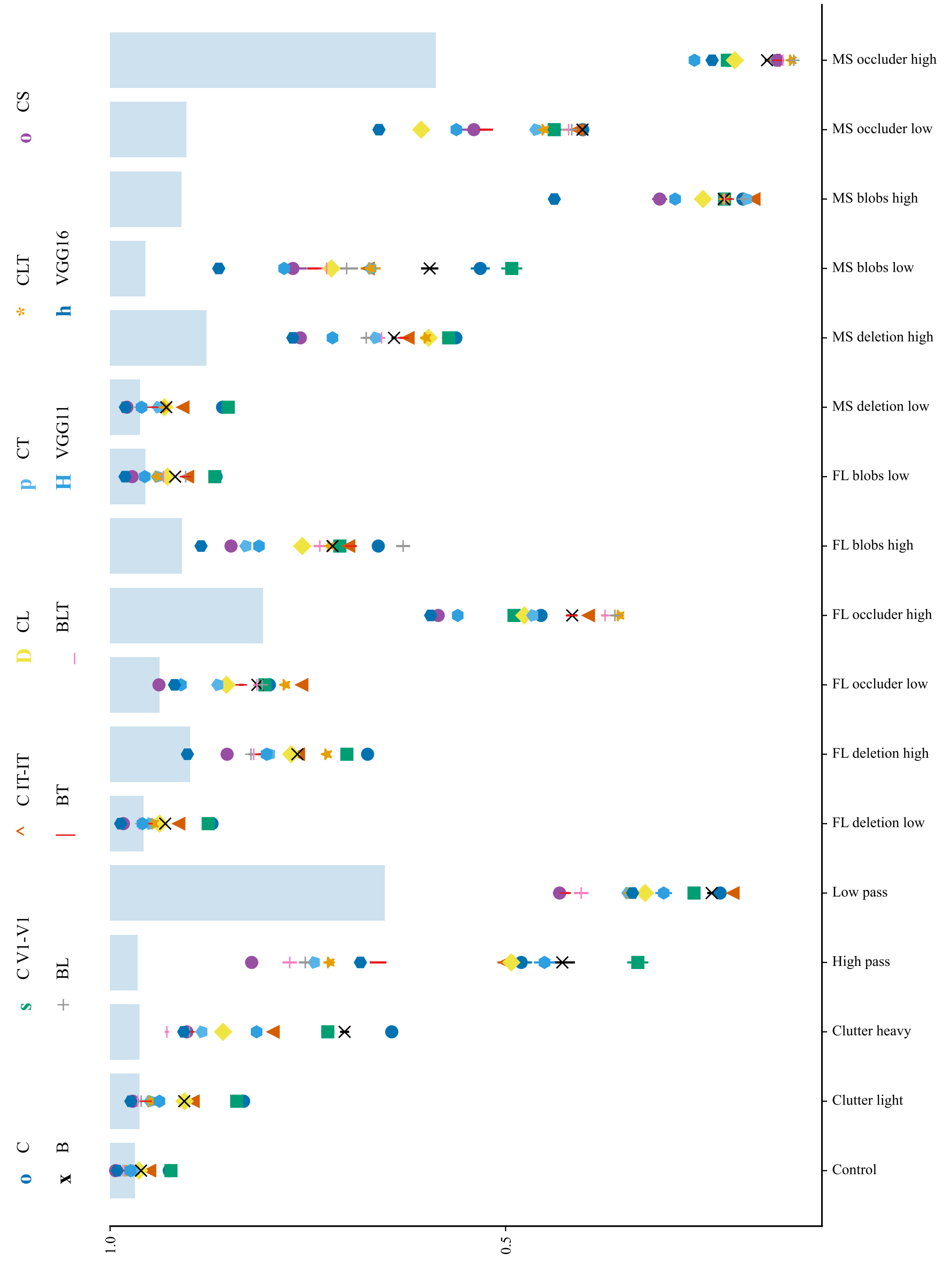


Figure S2: A less colour-heavy reproduction of figure 6. The colour palette has been adapted for more contrast. Shape distinctions have been added as well. *MS*: many small, *FL*: few large (referred to as *small* and *large* respectively in the main text).

##### 3.2 Confusion-based PCA plots

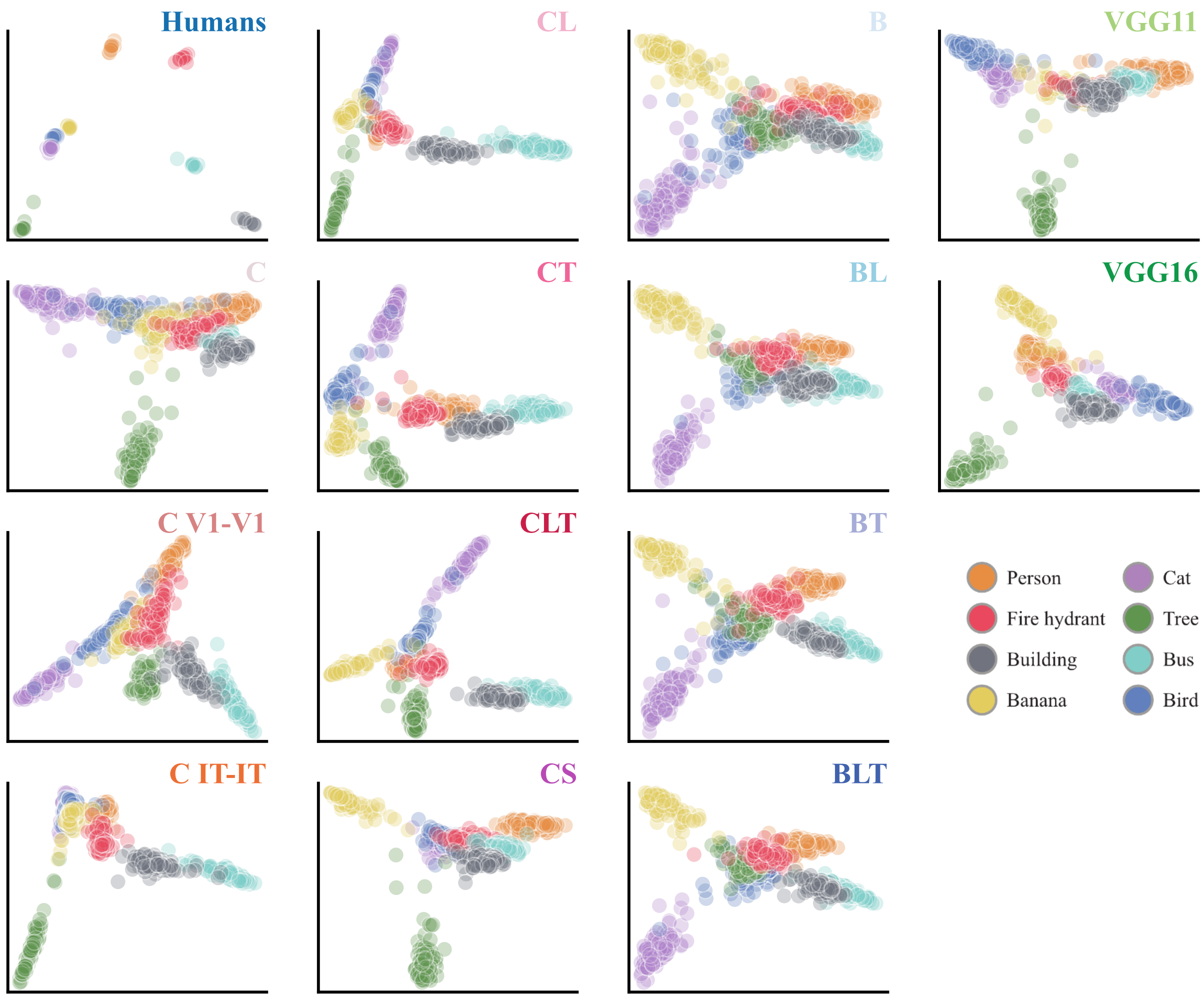


Figure S3: PCA plots for each model. Each dot represents an individual image, from one of eight possible categories (see legend at the bottom-right). For more clarity, graphs were aligned using General Procrustes Analysis (alignment to the average of all model graphs).

The PCA plots presented in figure S3 are derived from the image-wise confusion data. All models were presented with the same 800 images (100 images per category, 8 categories), passed through 16 challenging manipulations. Humans were presented with 80 images (10 per category), and their data was processed in a similar way. For each individual image, data was pooled from all condition and for each model. Confusion scores were calculated, reflecting how many times each model had classified the image in each of the eight categories. This resulted in 800 8-long vectors on which a 2-component PCA was fitted. The resulting two-dimensional values are plotted, per model and colour-coded per category, on figure S3.

To align the 2-dimensional PCA data of the 13 DNN plots with the human plot, a ***Generalized Procrustes Analysis*** (GPA) was run. GPA is a statistical technique used to align configurations of points in multidimensional space. It works by iteratively rotating, translating, and scaling the points of each PCA plot, with the goal of minimising the sum of squared distances between corresponding points of the reference plot and the other plots. It preserves the variation captured by the PCA components, while making comparisons between models and human participants easier. The GPA was ran on the model PCA data with the human PCA data as a reference. The latter was duplicated ten times to match with the size of the data to transform (which does not change the direction of the results, and gives more weight to the 80 initial data points of the human data set).

Overall, even after the GPA, no largely visible agreement seems to emerge from the plots. This is reflected in the disparity values, with the average point-wise disparity ranging from $0.47$ to $0.84$ (average: $0.68$). These numbers show how little qualitative agreement models have with humans in aligning their image-wise confusion data.

##### 3.3 Participant-level confusion matrix correlation

We calculated participant-level correlations with model confusion matrices, taking a more conservative approach to confirm the disadvantage of adding recurrent connections to DNNs. We built a confusion matrix per subject (n=218) and correlated it with the confusion matrix of each model. With the correlation of each individual participant with each model, we calculated a 95% confidence interval around the mean, which confirmed our results.

This approach confirms results from the main text: CS and recurrent versions of B all have average participant-level correlation scores outside of and lower than the CI boundaries of the correlation scores of their baseline counterparts (boundaries of $0.09$ and $0.08$ for C and B, respectively). Additionally, all recurrent versions of C and B show correlations lower than VGG 16 (all below the CI lower boundary of $0.13$). These values are striking when compared to model-model confusion matrix correlations: all models have a relatively higher average correlation with other models (Pearson's r, seed-level correlation with all other models in the range $0.24 - 0.94$, with average $0.73$).

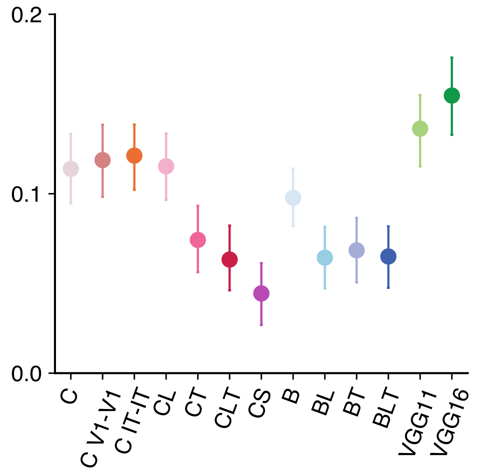


Figure S4: Average individual participant confusion matrix correlation with each model. Bars represent a 95% confidence interval around the mean. VGG 16 shows a higher correlation than any other model (all models below the lower CI boundary, $0.13$)

##### 3.4 Model-wise confusion matrix correlations

Supplementary figure S5 shows a matrix of correlations across models. Each square shows the average correlation value of 20 pairs of seed-level confusion matrices from two given models. Note the scale of the graph: correlations within models are mostly in the range of $0.3 - 1$. For reference, the confusion matrix correlations of models with humans range from $0.08$ to $0.43$. This indicates a larger agreement of models with each other than with human participants, sometimes in spite of architecture or model family. However, it is also noticeable that the range of within-model correlations sometimes overlaps with the range of correlations with humans. In particular, VGG16 seems to correlate better with humans (Pearson's r $0.43$) than with the smaller C models (e.g. with C: $0.24$, C V1-V1: $0.28$).

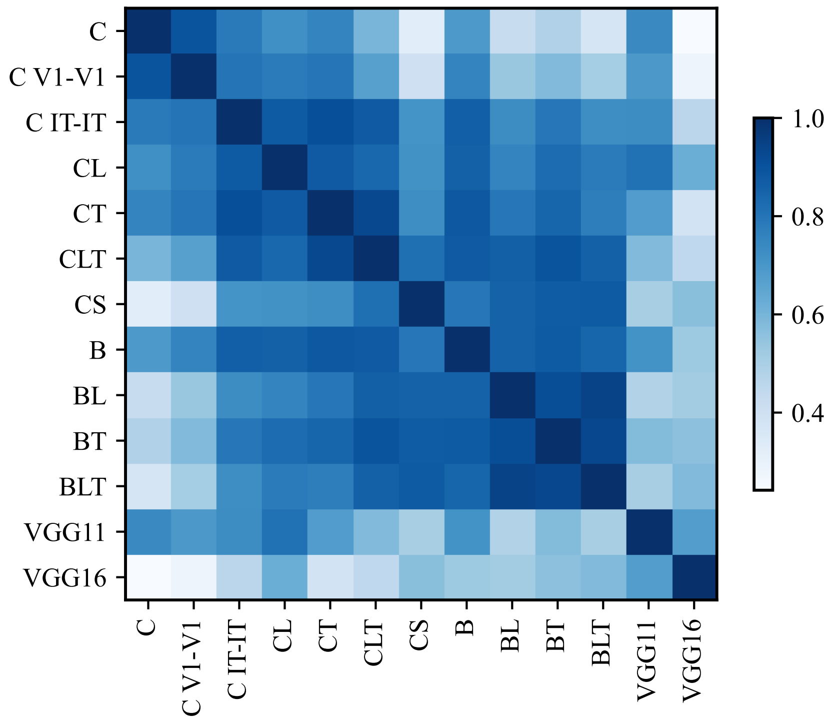


Figure S5: Correlation matrix: confusion matrices across models. Each cell represents the Pearson’s r correlation between the confusion matrices of two models.
